# IL-13 decreases susceptibility to airway epithelial SARS-CoV-2 infection but increases disease severity in vivo

**DOI:** 10.1101/2024.07.03.601941

**Authors:** Shreya Ghimire, Biyun Xue, Kun Li, Ryan M. Gannon, Christine L. Wohlford-Lenane, Andrew L. Thurman, Huiyu Gong, Grace C. Necker, Jian Zheng, David K. Meyerholz, Stanley Perlman, Paul B. McCray, Alejandro A. Pezzulo

## Abstract

Treatments available to prevent progression of virus-induced lung diseases, including coronavirus disease 2019 (COVID-19) are of limited benefit once respiratory failure occurs. The efficacy of approved and emerging cytokine signaling-modulating antibodies is variable and is affected by disease course and patient-specific inflammation patterns. Therefore, understanding the role of inflammation on the viral infectious cycle is critical for effective use of cytokine-modulating agents. We investigated the role of the type 2 cytokine IL-13 on SARS-CoV-2 binding/entry, replication, and host response in primary HAE cells in vitro and in a model of mouse-adapted SARS-CoV-2 infection in vivo. IL-13 protected airway epithelial cells from SARS-CoV-2 infection in vitro by decreasing the abundance of ACE2- expressing ciliated cells rather than by neutralization in the airway surface liquid or by interferon-mediated antiviral effects. In contrast, IL-13 worsened disease severity in mice; the effects were mediated by eicosanoid signaling and were abolished in mice deficient in the phospholipase A2 enzyme PLA2G2D. We conclude that IL-13-induced inflammation differentially affects multiple steps of COVID-19 pathogenesis. IL-13-induced inflammation may be protective against initial SARS-CoV-2 airway epithelial infection; however, it enhances disease progression in vivo. Blockade of IL-13 and/or eicosanoid signaling may be protective against progression to severe respiratory virus-induced lung disease.

**RESEARCH IN CONTEXT:** *Evidence before this study:* Prior to this study, various pieces of evidence indicated the significant role of cytokines in the pathogenesis and progression of COVID-19. Severe COVID-19 cases were marked by cytokine storm syndrome, leading to immune activation and hyperinflammation. Treatments aimed at modulating cytokine signaling, such as IL-6 receptor antagonists, had shown moderate effects in managing severe COVID-19 cases. Studies also revealed an excessive production of type 2 cytokines, particularly IL-13 and IL-4, in the plasma and lungs of COVID-19 patients, which was associated with adverse outcomes. Treatment with anti-IL-13 monoclonal antibodies improved survival following SARS-CoV-2 infection, suggesting that IL-13 plays a role in disease severity. Type 2 cytokines were observed to potentially suppress type 1 responses, essential for viral clearance, and imbalances between these cytokine types were linked to negative COVID-19 outcomes. These findings highlighted the complex interactions between cytokines and the immune response during viral infections, underscoring the importance of understanding IL-13’s role in COVID-19 and related lung diseases for developing effective therapeutic interventions.

*Added value of this study:* In this study, we explored the impact of IL-13-induced inflammation on various stages of the SARS-CoV-2 infection cycle using both murine (in vivo) and primary human airway epithelial (in vitro) culture models. Our findings indicated that IL-13 provided protection to airway epithelial cells against SARS-CoV-2 infection in vitro, partly by reducing the number of ACE2- expressing ciliated cells. Conversely, IL-13 exacerbated the severity of SARS2-N501Y_MA30_-induced disease in mice, primarily through Pla2g2d-mediated eicosanoid biosynthesis.

*Implications of the available evidence:* Current evidence indicates that PLA_2_G2D plays a crucial role in the IL-13-driven exacerbation of COVID-19 in mice, suggesting that targeting the IL-13-PLA2G2D axis could help protect against SARS-CoV-2 infection. These insights are important for clinical research, especially for studies focusing on drugs that modify IL-13 signaling or modulate eicosanoids in the treatment of asthma and respiratory virus-induced lung diseases.

## INTRODUCTION

The implementation of safe and effective vaccination made COVID-19 largely preventable and changed the course of the pandemic. Moreover, selected treatments with modest efficacy have been approved for people recently diagnosed with COVID-19. The combination of antiviral drugs nirmatrelvir and ritonavir, known as Paxlovid, can decrease the risk of progression when given early after diagnosis (1). Severe COVID-19 is characterized by cytokine storm syndrome and immune activation; therefore, promising treatments aimed at modulating cytokine signaling have been developed. For example, suppression of interleukin 6 (IL-6) signaling with receptor antagonists have moderate effects in severe cases (2). However, we continue to lack effective therapies to stop disease progression once respiratory failure ensues in COVID-19 and other respiratory virus-induced diseases.

Other cytokine targeting strategies have also shown promise. Several observations suggest that the type 2 cytokine, interleukin 13 (IL-13) which is associated with asthma, chronic bronchitis, and allergies, may be related to the severity of COVID-19: 1) In the plasma and lungs of COVID-19 patients, IL-13 and IL-4 were produced in excess (3); 2) treatment with an anti-IL-13 monoclonal antibody improved survival following SARS-CoV-2 infection (4) and is linked to a lower risk of mortality and improved lung function in humans (5, 6); and 3) Type 2 cytokines, like IL-4 and IL-13, may suppress type 1 responses necessary for viral clearance (7–12) and type 2/type 1 imbalances are linked to negative COVID-19 outcomes (13).

Understanding the role of inflammation on the outcomes of viral infection is important for therapeutic drug development and for selecting the right time and patients for the treatment. For example, the use of corticosteroids such as dexamethasone, compared with usual care or placebo, reduced mortality rate among hospitalized COVID-19 patients receiving mechanical ventilation; however, this effect did not extend to patients who did not require respiratory support (14, 15). Similarly, treating hospitalized COVID-19 patients with remdesivir, an antiviral drug (16), or baricitinib, an oral selective Janus kinase 1/2 inhibitor with anti-inflammatory properties, shortened recovery time and improved clinical status, notably for those on high-flow oxygen or ventilation (17).

Overall, previous studies suggest that type 2 inflammation in people with chronic lung disease has complex effects on COVID-19 progression. A better understanding of the role of IL-13 at each step of viral pathogenesis may facilitate development off effective treatments to prevent disease progression. In this study, we hypothesized that IL-13 affects susceptibility to SARS-CoV-2 infection and systematically investigated the role of IL-13 induced inflammation on different steps along the infectious cycle of SARS-CoV-2 and virus-induced lung disease.

## RESULTS

### IL-13 induced inflammation protects human airway epithelia from SARS-CoV-2 infection in vitro

Previous studies showed that type 2 airway inflammation decreases expression of the SARS-CoV-2 receptor ACE2 (Angiotensin Converting Enzyme 2) but increases expression of the entry factor TMPRSS2 (Transmembrane serine protease 2) in airway epithelia (18). In contrast, the pleiotropic cytokine TNFα and the type 17 cytokine IL-17 may have the opposite effect in human cells (19, 20). We hypothesized that IL-13-induced inflammation would protect primary human airway epithelia (HAE) from SARS-CoV-2 infection, whereas IL-17 with TNFα would synergistically enhance susceptibility to the virus. We pre-treated HAE cultured at the air-liquid interface with 20 ng/mL recombinant IL-13 or 20 ng/mL recombinant IL-17 combined with 10 ng/mL tumor necrosis factor alpha (TNFα) for 4 or 55 days. The cytokines were applied to both apical and basolateral cell surfaces. We then infected HAE with SARS-CoV-2 (MOI 0.1) and analyzed the cells 6 or 72 hours post-infection **(Fig. 1a)**. We measured viral titers in apical washes and used flow cytometry to enumerate cells positive for the nucleocapsid protein (N-protein) 72 hours post-infection (hpi). We found that IL-13, but not IL-17+TNFα exposed cells, were protected from SARS-CoV-2 infection when treated for 55 days. These data suggest that IL-13-induced epithelial remodeling (21) protects human airway epithelial cells from SARS-CoV-2 **(Fig. 1 b, c)**.

**Figure 1.**
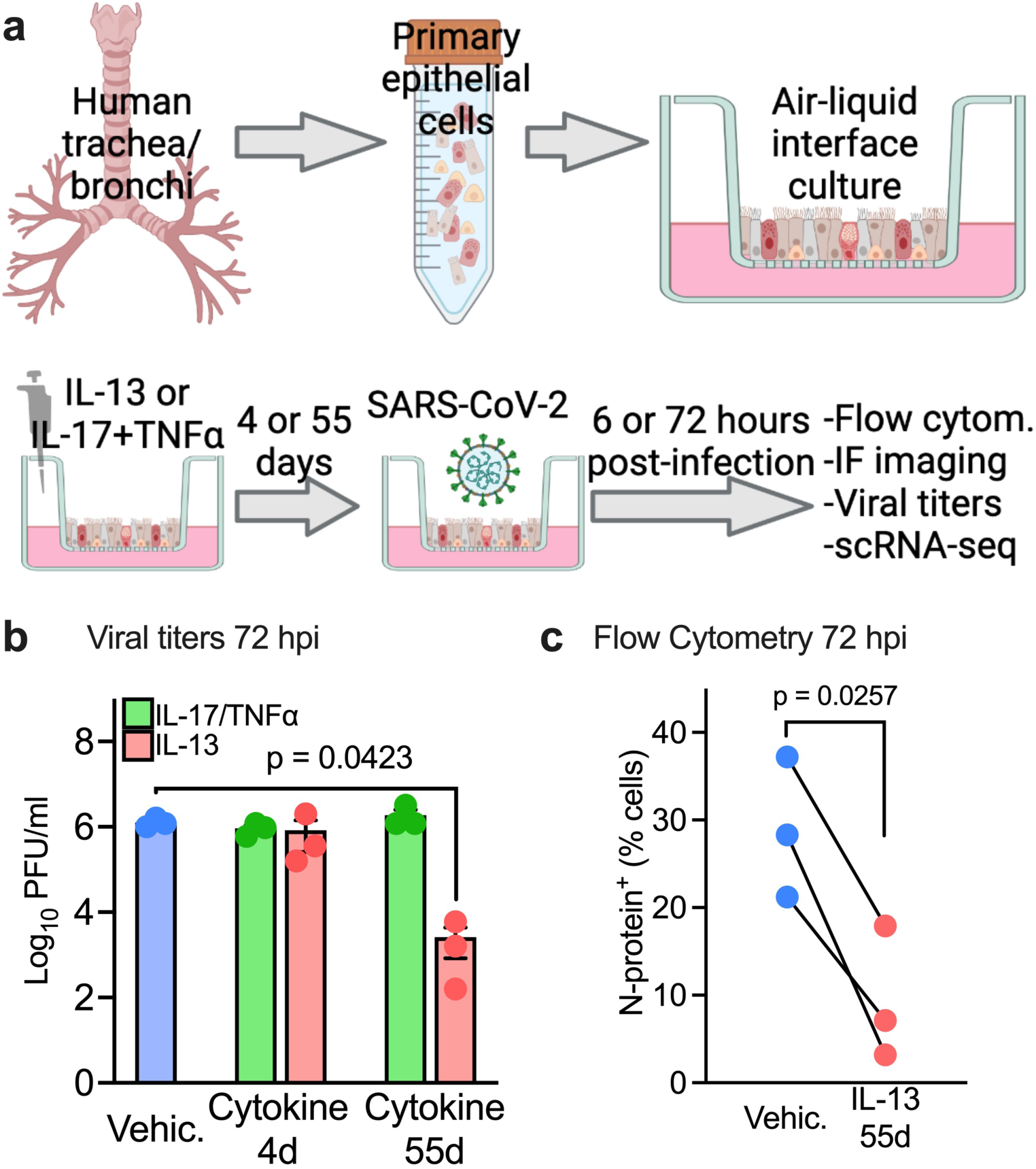
IL-13 induced inflammation protects human airway epithelia from SARS-CoV-2 infection in vitro. (a) Experimental conditions/in-vitro model: IL-13 (20ng/mL) or IL-17 (20ng/mL) +TNFα (10ng/mL) were added to well-differentiated air-liquid interface (ALI) primary human airway epithelia (HAE) for 4 or 55 days prior to infection with SARS-CoV-2(MOI 0.1). Samples were analyzed at 6 or 72 hours post infection (hpi). (b) Viral titers/PFU at 72hpi. Data represent the mean ± SEM and p value for one-way ANOVA test. n =3 (c) Flow cytometry showing % of cells expressing N protein at 72hpi. n= epithelia from 3 human donors, p value shown for paired t-test. (a) was created with Biorender.com

### IL-13 induced goblet cell metaplasia (GCM) does not affect surface viral neutralization or viral replication in primary human airway epithelial cells

Our data show that IL-13 induced inflammation protects HAE from SARS-CoV-2 infection in vitro. Previous reports show that IL-13 can modulate innate immune factors in the airway surface liquid that can neutralize viruses and kill bacteria (22, 23). We hypothesized that airway surface liquid from epithelia with IL-13 induced goblet cell metaplasia neutralizes the virus to protect HAE from SARS-CoV-2. We collected airway surface liquid (ASL) from 6 vehicle- and IL-13-exposed (55 days) donor epithelia and performed a neutralization assay using Sendai virus (mouse parainfluenza virus type 1, previously shown to be neutralized by airway surface liquid (24)), SARS-CoV-2 pseudotyped VSV at various MOI, and SARS-CoV-2 (**Fig. 2**). We found that ASL from IL-13 exposed HAE neutralized Sendai virus significantly at MOI 5 and 10 with average neutralizing activity of 80% and 60%, respectively. In contrast, the neutralization of SARS-CoV-2 pseudotyped VSV and of SARS-CoV-2 by ASL from IL-13 treated HAE was less than 0.5% at all MOI tested. Because the neutralizing activity of IL-13 treated HAE was minimal, it is unlikely that IL-13 induced changes in secreted antiviral factors can explain the protection from SARS-CoV-2 infection after IL-13.

**Figure 2.**
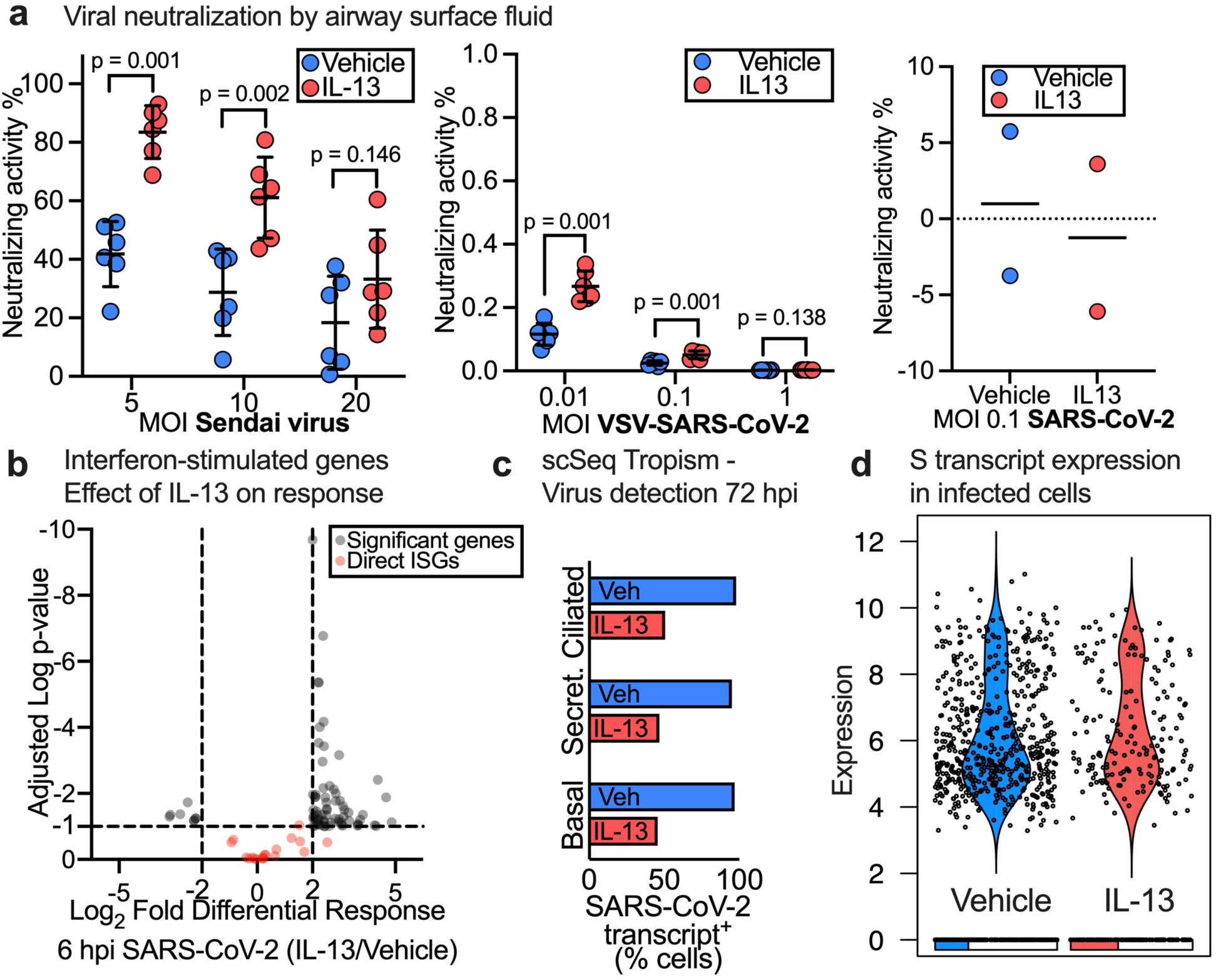
IL-13 induced goblet cell metaplasia (GCM) does not affect viral neutralization at the airway surface or viral replication. (a) Neutralization of Sendai virus (left), SARS-CoV-2 pseudo typed VSV (middle) and WT-SARS-CoV-2 (right) by airway surface liquid (ASL) collected from Vehicle and IL-13 treated (20ng/mL) HAE. n= epithelia from 6 or 2 humans, p-value shown for Student’s unpaired *t* test and data represent mean ± SD. (b) Volcano plot for IL-13 (20ng/mL, 55 days) vs Vehicle pre-treated ciliated cells at 6 hpi with SARS-CoV-2. Dotted line represents cutoffs for significant genes (log2FC ± 2, adj p-value=0.1). Significant genes and ISGs directly affected by virus shown. (c) Effect of IL-13 treatment (20ng/mL, 55 days) on detection of SARS-CoV-2 transcript-positive cells in each major epithelial cell type. (d) Violin plot of S transcript in Vehicle vs IL-13 (20ng/mL, 55 days) treated HAE at 72 hpi with SARS-CoV-2. Each dot represents a SARS-CoV-2 transcript-positive cell and the bar at the bottom shows the proportion of cells with zero expression in each group.

We next hypothesized that IL-13 affected expression of the subset of interferon stimulated genes that directly impact viral replication (25). We measured the differential response of IL-13- vs. vehicle-pre-treated cells to SARS-CoV-2 at 6 hpi using single cell RNA-seq (**Supplementary FiguresS1 and S2**) and mixed model analysis (described in methods); we focused on ciliated cells because they are the initial target for infection by SARS-CoV-2 (26). We found that IL-13-treated HAE responded to SARS-CoV-2 differentially at 6 hpi as shown by the number of significant genes in **Fig. 2b**, however, we found that IL-13 did not affect the expression of interferon-stimulated genes (ISGs) that directly inhibit viral replication (25).

To determine the effect of IL-13 on susceptibility of epithelial cell types to infection and on viral replication in infected cells, we measured detection of SARS-CoV-2 transcript-positive cells in each major epithelial cell type and viral transcript levels in cells 72 hours post exposure to SARS-CoV-2. We used a stringent method to account for background scRNA-seq library contamination (**Supplementary FigureS3**). We found that IL-13 decreased the abundance of SARS-CoV-2 transcript-positive cells in all major epithelial cell types (**Fig. 2c**); however, the average levels of SARS-CoV-2 transcripts in infected cells did not differ between vehicle- and IL-13-treated epithelia (**Fig. 2d and Supplementary FigureS4)**. Using mixed model analysis, we identified genes induced by 72 hours of SARS-CoV-2 exposure in vehicle and IL-13 (4days) treated infected ciliated cells. We also characterized the differential response of IL-13 treatment in infected ciliated cells exposed to SARS-CoV-2 for 72 hours (**Supplementary FigureS5, Supplementary Table1**). In summary, our results show that IL-13 induced GCM affects neither viral neutralization by ASL nor viral replication and suggest that IL-13 affects either viral-epithelial binding or cellular entry.

### IL-13 induced GCM protects HAE from SARS-CoV-2 in vitro by decreasing the abundance of receptor-expressing cells

Previous studies have shown that type 2 inflammation affects the expression of SARS-CoV-2 entry factors ACE2 and TMPRSS2 in airway epithelia (27). We hypothesized that IL-13 induced GCM protects HAE from SARS-CoV-2 infection by decreasing the abundance of receptor-expressing cells, thus limiting viral-epithelial binding or cellular entry. Using our single cell RNA-seq data, we measured the proportion of major cell types in uninfected vehicle- and IL-13-treated (55 days) samples. We found that compared to vehicle, IL-13 treated samples had significantly fewer secretory and ciliated cells, and a greater abundance of basal and goblet cells (**Fig. 3a**). Next, we measured the transcript expression of the viral entry factors ACE2, TMPRSS2, and furin in each of those major cell types from vehicle and IL-13 (55 days) treated groups. We found that IL-13 did not affect ACE2 expression in ciliated cells. IL-13 did decrease ACE2 expression in basal cells, although basal cells are not considered primary viral infection targets. The expression of TMPRSS2 was higher in IL-13-treated secretory cells, whereas FURIN expression was unaffected (**Fig. 3b**). Overall, we found that while IL-13 does not affect the expression of ACE2 in ciliated cells, it does decrease the overall abundance of ciliated cells.

**Figure 3.**
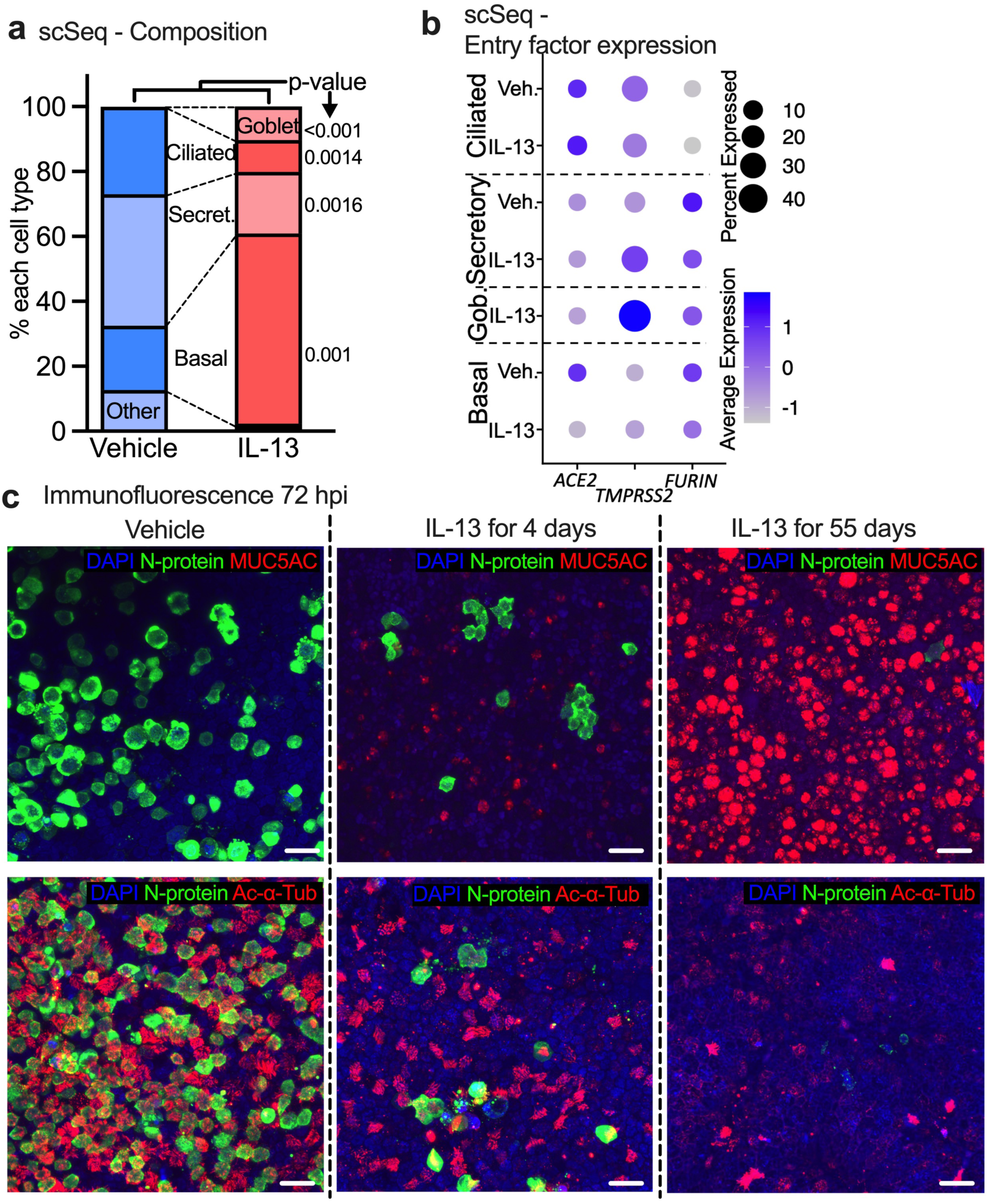
IL-13 induced GCM protects HAE from SARS-CoV-2 in vitro by decreasing the abundance of receptor-expressing cells. (a) Proportion of common cell types in vehicle- and IL-13-treated (20ng/mL, 55 days) samples (no virus exposure) determined by single cell RNA-seq data. n=3, p-value shown for unpaired t test. (b) Dot plot showing the effect of IL-13 treatment (20ng/mL, 55 days) on viral entry factor expression in uninfected HAE. Size of the dot represents % of the cell expressing the gene and color scale shows the average expression level. n = 3 donors (c) Representative image of immunofluorescence confocal microscopy for DAPI (nuclei, blue), N-protein (virus, green), and MUC5AC (goblet cells, red, top) or acetylated alpha tubulin (ciliated cells, red, bottom).

To further determine the effects of IL-13 on SARS-CoV-2 infection of HAE, we performed confocal microscopy for DAPI (nuclei), N-protein (virus), MUC5AC (goblet cells), and acetylated alpha tubulin (Ac-α-Tub, ciliated cells) in Vehicle and IL-13-treated (4 days, 55 days) HAE at 72 hpi with SARS-CoV-2. A representative image for each condition is shown in **Fig. 3c**. We found that epithelia treated with IL-13 for 55 days had 1) a significant increase in goblet cells, shown by MUC5AC (red, top row), 2) a significant decrease in the abundance of ciliated cells, shown by Ac-α-Tub (red, bottom row) and, 3) a significant decrease in SARS-CoV-2 virus antigen, shown by N-protein (green). These results from confocal microscopy match the quantification obtained from scRNA-seq data and the quantification of infection by titers and N-protein in Figure 1B, C. Moreover, immunostaining for ACE2 protein (**Supplementary FigureS6**) supported the scRNA-seq finding that IL-13 decreases the abundance of ACE2^+^ ciliated cells but not the abundance of ACE2 in each cell. In summary, our findings suggest that IL-13-induced GCM protects airway epithelia from SARS-CoV-2 infection in vitro, in part by decreasing the abundance of ciliated cells which are one of the main targets for SARS-CoV-2 tropism.

### IL-13 enhances SARS2-N501Y_MA30_-induced disease in vivo

IL-13 can induce airway inflammation including goblet cell metaplasia, sub-epithelial fibrosis, and immune cell recruitment; however, it also possesses anti-inflammatory properties. The anti-inflammatory properties of IL-13 include the polarization and activation of immunosuppressive M2 (arginase- and macrophage mannose receptor - CD206-positive) macrophages and the differentiation of regulatory T cells, which are compromised in severe COVID-19 patients (28–30). We hypothesized that IL-13-induced dysfunction of immune responses contributes to increased susceptibility to COVID-19. Therefore, we infected young (6-10 weeks old) BALB/c mice with 1,000 PFUs of SARS2-N501Y_MA30_, a mouse adapted SARS-CoV-2 (31), after a pretreatment of 2.5 μg/day intranasal IL-13 (previously shown to induce lower airway goblet cell metaplasia in (21)) or Vehicle for 4 days (**Fig. 4**).

**Figure 4.**
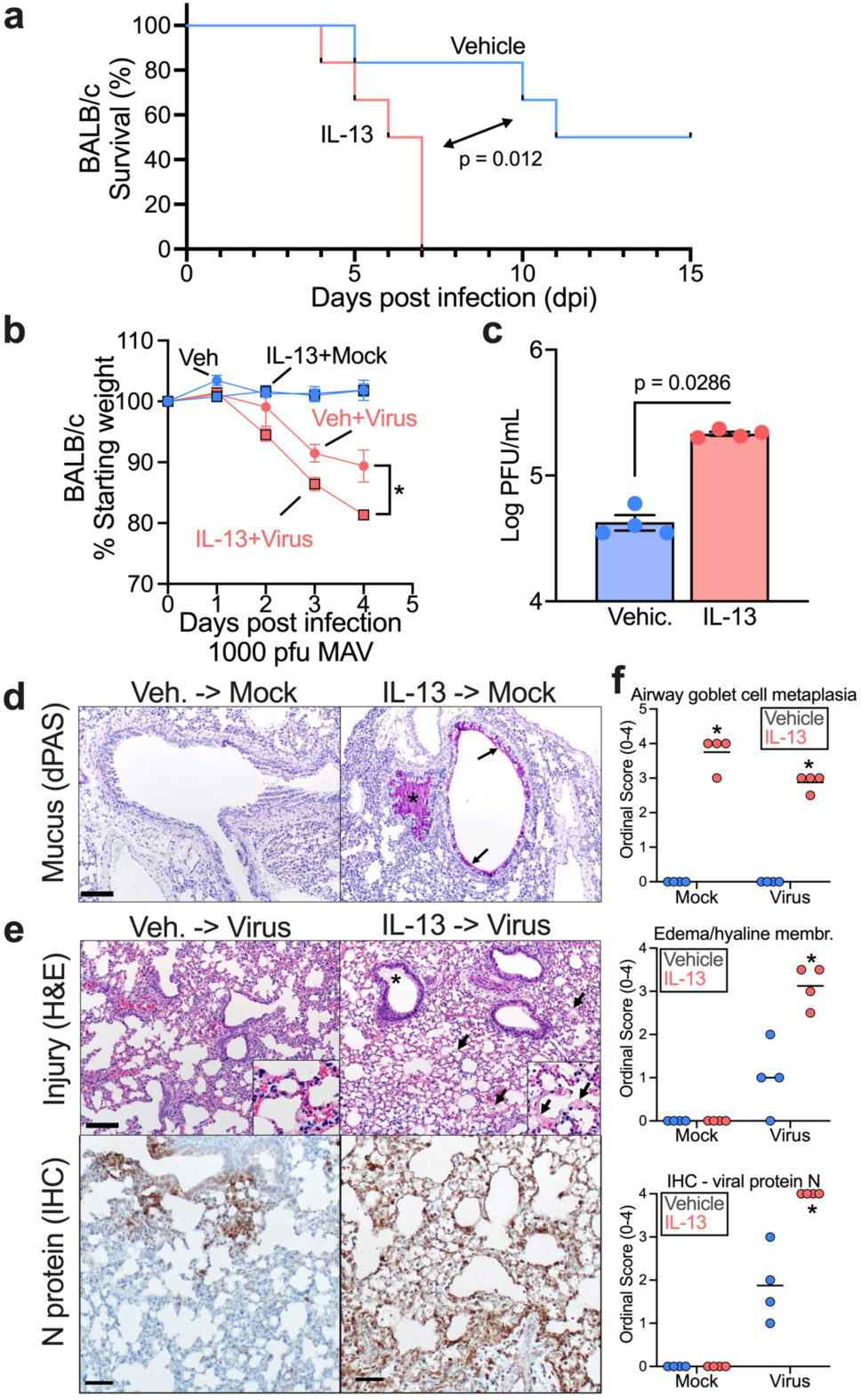
IL-13 enhances SARS2-N501Y_MA30_-induced disease in vivo. 6-10 week-old BALB/c mice were intranasally infected by 1,000 PFU of SARS2-N501Y_MA30_ after a 4-day pretreatment of 2.5 μg/day intranasal IL-13 or vehicle. (a) Survival rate and (b) weight were monitored daily. n = 4-6 mice for each group; mean ± SEM; for % starting weight of Veh + Virus vs IL-13+Virus, *p˂0.05; p-values in weight change and survival curve are Student’s unpaired *t* test and log-rank (Mantel-Cox) tests, respectively. (c) Quantification of virus loads at 5 dpi in homogenized mouse lungs by plaque assay using VeroE6 cells. n = lung tissue from 4 mice; p value: Student’s unpaired *t* test. (d) Representative sections from uninfected mouse lungs treated with vehicle or IL-13treatment and stained with diastase-pretreated periodic acid-Schiff (dPAS). Note the increased dPAS+ staining of the epithelium (arrows) and within the airway (asterisk), (e) Representative sections from infected mouse lungs with vehicle or IL-13 treatment and stained with H&E (top) or immunostained for SARS-CoV-2 N protein. Note the edema (asterisk, top right image) and hyaline membranes (arrows, top right image). Note the expansive N protein staining in the bottom right versus bottom left representative images. Histopathology staining from 4 mice/group. (f) Ordinal histopathological scoring of airway goblet cell metaplasia (mucus-producing goblet cells) (top panel), edema/hyaline membrane (middle panel), and SARS-CoV-2 N protein expression (bottom panel) in lungs from mock or infected mice. mean ± SEM; *p˂0.05; p value: Student’s unpaired *t* test

IL-13-treated mice had higher mortality and lost more weight upon infection compared to Vehicle (**Fig. 4a, b**). This result complements findings from a study showing that blockade of IL-13 with an antibody decreases SARS-CoV-2-induced mortality (4). To assess viral load, we titered virus in lung tissue from IL-13- or Vehicle-pretreated mice on day 5 (**Fig. 4c**). Consistent with faster disease progression, elevated viral loads were present in lung tissue of IL-13-pretreated mice. Histopathological analyses of tissue from mock infected mice and infected mice at day 5 post infection revealed evident goblet cell metaplasia and lung injury in IL-13-pretreated mice, with hypersecretion of mucus and evidence of acute lung injury (pulmonary edema/hyaline membranes) (**Fig. 4d, f**). Next, we performed bulk RNA-seq on lung tissue from SARS2-N501Y_MA30_ infected and uninfected mice, pretreated with either IL-13 or Vehicle. We conducted differential gene expression (DGE) analysis and pathway analysis to identify the effects of IL-13 on the response to SARS2-N501Y_MA30_. Pathway analysis using the GO: Biological process pathway database revealed downregulation of antigen processing and presentation of endogenous peptide antigen genes in IL-13-treated mice. Some other downregulated pathways in IL-13-treated mice were complement activation, cytolysis, xenobiotic catabolic and xenobiotic metabolic processes. (**Supplementary Figure S7, Supplementary Table2**). Qualitative histopathological analysis shows that samples from vehicle-treated mice developed regions of consolidation with atelectasis, increased cellularity by leukocytes and cellular debris with edema. In contrast, samples from IL-13-treated mice lacked regions of consolidation, had increased edema and hyaline membranes **(Supplementary FigureS8)**. In addition, we performed immunohistochemical detection of AIF1/IBA1 which is expressed by M2 macrophages. We found that samples from the IL-13-treated mice had an increase in AIF1^+^ cells; the AIF1+ cells were predominantly peri-venous in samples from mock-infected animals but were more diffusely localized with virus infection **(Supplementary FigureS9).**

These findings may explain the increased viral titer in IL-13-treated mouse lungs compared to the vehicle group. And suggest that pre-existing IL-13-induced inflammation may impair viral clearance or affects infection kinetics in addition to shifting the immune and epithelial response to infection.

### IL-13 induces dysregulation of eicosanoids in vivo

Our data reveal a discrepancy between the effects of IL-13 in airway epithelia in vitro (with only epithelial cells present) and in mice in vivo (epithelial, immune, mesenchymal cells, etc.). We hypothesized that the effects of IL-13 would vary between human airway epithelia in vitro and murine lungs in vivo. We carried out transcriptional profiling of primary airway epithelial cells (21-day IL-13/Vehicle treatment) and lung tissues of non-infected mice (4-day IL-13/Vehicle treatment) by bulk RNA-seq and then identified and compared differentially expressed genes (DEGs) between the IL-13 and vehicle groups (**Fig. 5a**). This analysis revealed that 457 and 1361 genes were differentially expressed at a p˂0.05 between IL-13 and Vehicle-treated primary airway epithelial cells and mice lungs, respectively. Whereas some genes responded similarly in vitro and in vivo (a selected sub-set of genes known to participate in IL-13-induced inflammation is shown in **Fig. 5a**), there were many differences between the groups.

**Figure 5.**
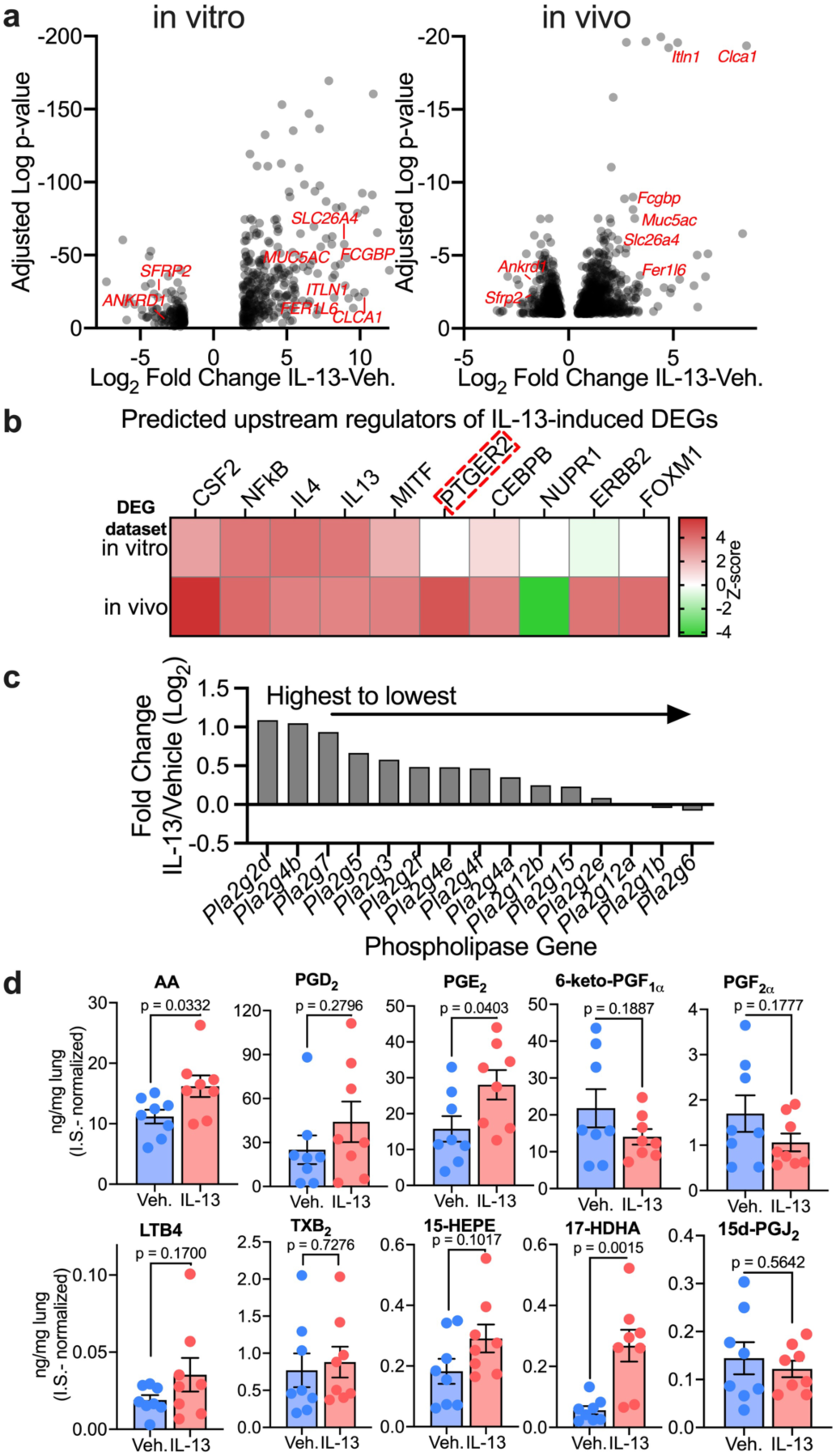
IL-13 induces dysregulation of eicosanoids in vivo. (a) RNA-seq analysis of IL-13 (20ng/mL)- or vehicle-treated primary human airway epithelial cells (21-day IL-13/Vehicle treatment, in vitro) and lungs of 6-10 week-old BALB/c mice (4-day, 2.5 μg/day intranasal IL-13/Vehicle treatment, in vivo). Volcano plots of differential gene expression analysis in vitro and in vivo of the effect of IL-13 compared to vehicle. n = epithelia from 8 humans (left); n = lung tissue from 4 mice (right). Selected genes relevant to goblet cell metaplasia are labeled in red. (b) Ingenuity Upstream Regulator Analysis of DEGs in IL-13- or vehicle-treated samples in vitro and in vivo. Predicted upstream regulators are sorted by absolute Z-score. *PTGER2* is highlighted as the top predicted regulator activated in vivo but not in vitro. (c) List of top 15 eicosanoids biosynthesis-associated phospholipase genes ranked from highest to lowest magnitude of IL-13 response. (d) Concentrations of prostaglandins in lungs of 8 week-old BALB/c mice treated with IL-13 or vehicle measured by LC-MS/MS calculated according to standard curves and normalized to internal standards (0.1 ng/μL for each I.S.). n = lung sample from 8 mice; mean ± SEM; p value is shown from Student’s unpaired *t* test.

We hypothesized that the discrepancy between the effect of IL-13 in vivo in murine lungs vs. in vitro in human airway epithelia was due to factors missing from the in vitro model; this may include a non-epithelial cell type or processes that require non-epithelial cell participation. To identify candidate drivers of IL-13-enhanced SARS2-N501Y_MA30_-induced disease, we conducted a comparative upstream regulator analysis (Ingenuity Pathway Analysis) of the response to IL-13 between in vitro epithelial cells and in vivo murine lungs. The data showed that both in vitro and in vivo, the top predicted upstream regulators of the effects of IL-13 on the transcriptome included both IL-4, IL-13 as expected, and other targets of the IL-13 signaling pathway including CSF2 and NFkB (**Fig. 5b**) (32, 33). These data validate the use of Upstream Regulator Analysis to predict drivers of transcriptional responses in the lungs. Next, we examined upstream regulators predicted as differentially activated in vitro and in vivo. The results suggest that *PTGER2* is activated by IL-13 in vivo, but not in vitro (**Fig. 5b**). *PTGER2* is a receptor of prostaglandin E2, highly expressed in kidney, lung, and the central nervous system, playing an important role in the resolution of inflammation and downregulation of immune responses (34, 35). Moreover, multiple eicosanoid biosynthesis-associated phospholipase genes were upregulated in response to IL-13, suggesting eicosanoids could mediate the observed IL-13-induced enhancement of COVID-19 (**Fig. 5c**).

Whereas eicosanoid biosynthesis can be transcriptionally regulated, control of eicosanoid levels and function is complex and multifactorial; eicosanoid pathway transcript and effector molecule abundance may correlate in some contexts but not others. We analyzed lung tissue from 8-week-old BALB/c mice treated with vehicle or IL-13 using LC-MS/MS to determine whether IL-13 alters lung tissue eicosanoid abundance. LC-MS/MS analysis confirmed induction of the eicosanoid precursor arachidonic acid (AA), prostaglandin E2 (PGE_2_), and the specialized pro-resolvin mediator precursor 17(S)-hydroxy docosahexaenoic acid (17(S)-HDHA) in the lungs of non-infected mice treated with IL-13 (**Fig. 5d**). In humans with mild and severe COVID-19, accumulation of various prostaglandins has also been reported in the serum (36) and BAL (37, 38). In sum, these data predict that the divergent effects of IL-13 on SARS-CoV-2 infection in vitro and in vivo are due to dysregulation of eicosanoids.

### IL-13 enhances SARS2-N501Y_MA30_-induced disease in vivo via eicosanoid dysregulation

We decided to study the role of the secreted Phospholipase A2 Group IID (*Pla2g2d*) on IL-13-enhanced SARS2-N501Y_MA30_ disease for three main reasons: 1) phospholipase abundance is the first and a rate-limiting step in eicosanoid function (39), 2) *Pla2g2d* was the top phospholipase induced by IL-13 in murine lungs, and 3) we previously showed that *PLA2G2D* is an important determinant of SARS-CoV (MA15, mouse adapted SARS-CoV) and SARS2-N501Y_MA30_-induced disease in mice and its effects are age-dependent (31, 40). We hypothesized that *Pla2g2d* is required for IL-13-enhanced SARS2-N501Y_MA30_ disease in middle-aged C57BL/6J mice.

We infected 8-9 month old wild-type and *Pla2g2d*^-/-^ C57BL/6J mice with 1,000 PFU of SARS2-N501Y_MA30_ (lethal dose). Similar to our findings in young BALB/c mice, we found increased SARS2-N501Y_MA30_-induced mortality in mice pretreated with IL-13 relative to vehicle. However, the *Pla2g2d*^-/-^ mice were protected from IL-13-enhanced SARS2-N501Y_MA30_-induced disease, with similar survival rates between IL-13- and Vehicle-treated mice (**Fig. 6a**). In both wild-type and *Pla2g2d*^-/-^ mice, IL-13 pre-treatment increased virus titers on day 2 post infection, however, this effect was reduced in *Pla2g2d*^-/-^ mice, implying the effect of IL-13 on virus load is partially dependent on eicosanoids (**Fig. 6b**). Overall, these results suggest that IL-13 enhances viral load and worsens the outcome of SARS2-N501Y_MA30_-induced murine disease through *Pla2g2d*-mediated eicosanoid biosynthesis.

**Figure 6.**
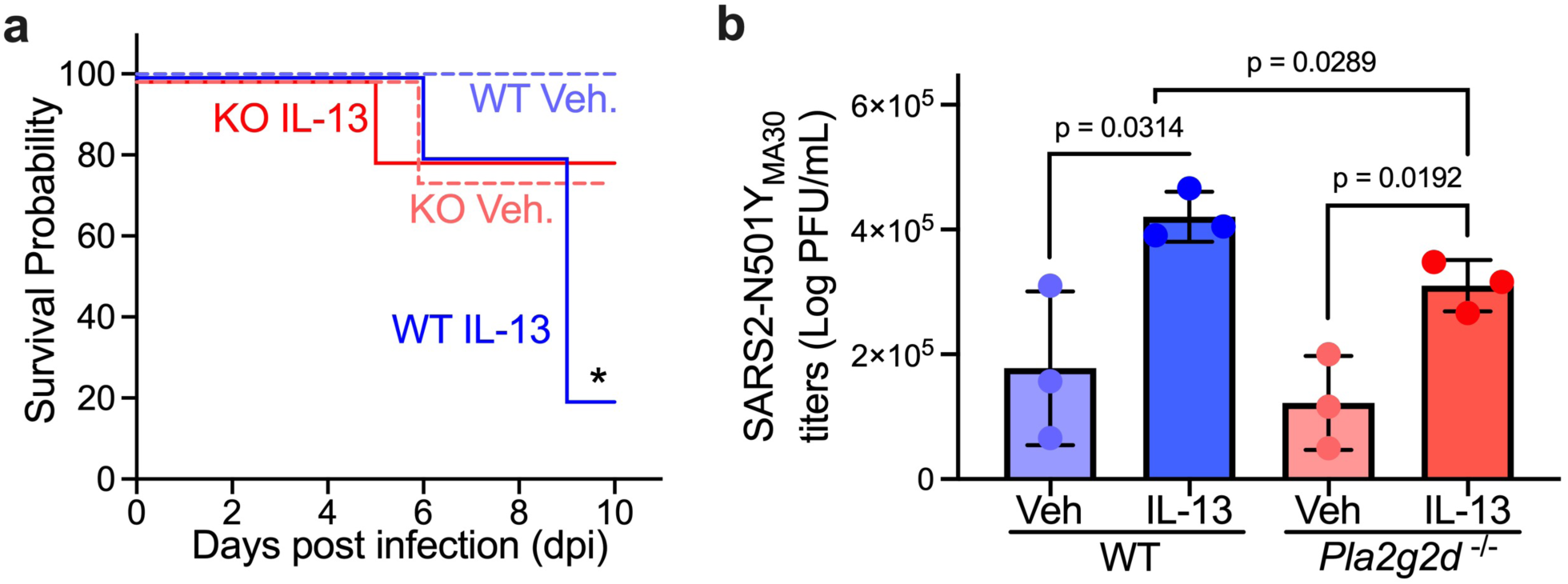
IL-13 enhances SARS2-N501Y_MA30_-induced disease in vivo via eicosanoid dysregulation. 8-9 month-old C57BL/6J wild-type (WT) and PLA2G2D-knock-out (KO) mice were pretreated with IL-13 and vehicle for 4 days and infected with 1,000 PFU SARS2-N501Y_MA30_ on day 5. (a) Survival curve of infected wild type and PLA2G2D-knock-out mice. n = 4-5 mice for each group; mean ± SEM; *p˂0.05; p value is shown from log-rank (Mantel-Cox) tests for WT Vehicle vs WT IL-13. (b) Viral loads in lungs of infected WT and KO mice on day 2 post infection were assessed by plaque assay. n = lung sample from 3 mice; mean ± SEM; p value is shown from Student’s unpaired *t* test.

## DISCUSSION

Here, we investigated the effects of IL-13 induced inflammation on multiple steps of the SARS-CoV-2 infectious cycle using both in vitro and in vivo models. Moreover, we generated an atlas of the effects of IL-13- and TNFα/IL-17-induced inflammation on the time course of cellular responses of air-liquid interface primary airway epithelia to SARS-CoV-2 at the single cell level; this dataset contains 48 samples from three human subjects and will be useful to the scientific community. We found that IL-13 protected airway epithelial cells from SARS-CoV-2 infection in vitro in part by decreasing the abundance of ACE2- expressing ciliated cells. However, IL-13 worsened the outcome of SARS2-N501Y_MA30_-induced murine disease through *Pla2g2d*-mediated eicosanoid biosynthesis.

Our data show a time-dependent IL-13-driven protective effect against SARS-CoV-2 in vitro. IL-13 induces airway goblet cell metaplasia characterized by an increase in mucus-secreting goblet cells and a reduction of ciliated cell abundance, as we and others have shown (21, 41, 42). Consistent with other studies, our results show an IL-13-induced decrease in ACE2 and TMPRSS2 expression (18); however, our single cell RNA-seq data demonstrate that the effects of IL-13 on virus receptors and virus entry factors may be predominantly driven by changes in cell composition rather than by receptor expression within each cell type (43–46). SARS-CoV-2 preferentially targets ciliated cells (26, 45), whereas MERS-CoV mainly infects goblet cells (43, 47); interestingly, IL-13 increases susceptibility to MERS-CoV (43). Thus, these findings indicate that IL-13-induced changes in cellular composition modulate airway susceptibility to coronaviruses in vitro.

We found a discrepancy between the effects of IL-13 on airway epithelia in vitro and in mice in vivo. These results complement a recent study demonstrating that IL-13 receptor blockade with dupilumab protects K18-hACE2 transgenic mice (developed by the authors (48)) from severe COVID-19 (4). To identify potential factors underlying the discrepancy between our in vivo and in vitro studies, we performed a comparative upstream regulator analysis between in vitro HAE cells and in vivo lung tissue bulk RNA-seq data. In addition to expected upstream regulators predicted to be activated both in vitro and in vivo (e.g., IL-4, IL-13, and NFκB signaling), multiple inflammation-associated factors were upregulated in response to IL-13 in mouse lung tissue but not in airway epithelia cells. This suggests that murine COVID-19 is likely enhanced by IL-13-induced dysregulated innate immune responses. Given our recent finding that PLA2G2D and prostaglandin D2 signaling modulate susceptibility to SARS-CoV-2 (31), we focused on the upstream regulator analysis prediction of IL-13-enhanced *PTGER2* signaling in murine COVID-19, which suggests eicosanoid signaling dysregulation.

Phospholipases, cyclooxygenases, and prostaglandin synthases regulate eicosanoid signaling; eicosanoids interact with their corresponding receptors which are expressed in a cell type-dependent manner (49). The accumulation and activation of eicosanoids in immune effectors including dendritic and mast cells, play an important role in regulating inflammation and immune responses through both paracrine and autocrine actions (49, 50). PLA_2_G2D (phospholipaseA2 group 2D) and one of its downstream eicosanoid receptors, prostaglandin D_2_ receptor (PTGDR), were found to modulate the progression of autoimmune disease and disease driven by various coronaviruses including mouse hepatitis virus (MHV), SARS-CoV and SARS-CoV-2, as previously shown by us and others (31, 40, 51–53). Our study reveals that PLA_2_G2D is essential for IL-13-enhanced murine COVID-19 disease enhancement, suggesting that IL-13-PLA_2_G2D axis may be targeted to protect individuals from SARS-CoV-2 infection.

However, eicosanoid signaling is complex: 1) phospholipase abundance can be a rate-limiting step, 2) the specific proteinoid synthesized is milieu-dependent and can be mediated by a single cell or a specific combination of cells, 3) the effects of receptor signaling are cell type-specific, and 4) there is significant ligand-receptor activity overlap between the different prostaglandins and their receptors. Moreover, the effects of prostaglandin signaling on virus-induced disease are likely time-dependent. Therefore, it will be essential to further dissect the role of IL-13 and eicosanoid signaling at different doses and time-points in SARS-CoV-2-induced disease.

There are some limitations to our approach. We studied only one variant of SARS-CoV-2; susceptibility to later variants may be affected differently by IL-13 or prostaglandins. In addition, we only investigated the effect of the type 2 cytokine IL-13 on SARS-CoV-2; the role of IL-13 varies in different inflammatory lung diseases, including in different asthma endotypes (54). To facilitate mechanistic interpretation, we focused on IL-13 as an inflammatory trigger, however, unlike allergen-sensitization asthma models, the IL-13 instillation murine model primarily affects innate rather than adaptive immunity. Previous studies have shown that whereas C57BL/6 mice show T_h_1- and M1-dominant immune response, BALB/c mice show more T_h_2- and M2 dominant immune responses (55, 56); strain-dependent differences could confound our in vivo infection model results, however, both young BALB/c and middle-aged C57BL/6 mice showed IL-13-enhanced disease. We also did not explore all possible mechanisms of virus-susceptibility modulation by IL-13 suggested by our transcriptomic analysis. Recent studies reported that IL-13 modulates hyaluronan polysaccharide and keratan sulfate abundance and influences the outcome of SARS-CoV-2 infection in vitro and in vivo (4, 41, 57).

The incidence of severe COVID-19 is fortunately steadily decreasing as a larger proportion of the population has been vaccinated and/or acquired immunity after infection. However, mechanistic research to better understand how type 2 inflammation affects susceptibility to severe SARS-CoV-2 is fundamental for multiple reasons, including: 1) future pandemic coronaviruses may behave similarly to SARS-CoV-2, 2) knowledge of the role of IL-13 on SARS-CoV-2 susceptibility may be relevant to other current respiratory viruses that cause respiratory failure, 3) understanding the role of currently approved therapies such as IL-4/IL-13 receptor antagonists, on coronavirus susceptibility is important for clinical decision-making during hospitalization of asthmatics, and 4) the efficacy of current therapies for patients with COVID-19 or other respiratory virus-induced respiratory failure is limited. Our findings have important implications for clinical studies including trials of IL-13 signaling-modifying or eicosanoid-modulating drugs in asthma and respiratory virus-induced lung disease (31, 58, 59).

## METHODS

### Differentiated primary airway culture

Primary epithelial cells were isolated from the human trachea and bronchi by enzymatic digestion and were grown at air-liquid interface using previously described method (60). Passage −0 primary differentiated human airway epithelia (HAE) from 3 human donors were obtained from the University of Iowa Cells and Tissue Core. Cell cultures were treated with 20 ng/mL recombinant IL-13 or 20 ng/mL recombinant IL-17 combined with 10 ng/mL tumor necrosis factor alpha (TNFα) for 4 or 55 days. The cytokines were applied to both apical and basolateral side of HAE. The cells were then infected with SARS-CoV-2 (MOI 0.1), the strain isolated from the first US patient (2019-nCoV/USA-WA1/2020, GenBank:MN985325.1) for 6 or 72 hr.

### Flow cytometry (neutralization assay)

For neutralizing assay, SeV-eGFP and VSV-eGFP-SARS-CoV-2 were pre-incubated with 100μL apical wash (AW) harvested from the surface of IL-13-pretreated or Vehicle (PBS)-pretreated ALI cultured HAE cells. Following a 2 hr incubation, the SeV-eGFP-AW and VSV-eGFP-SARS-CoV-2-AW mixtures were added to LLC-MK2 cells or VeroE6-TMPRSS2-ACE2 cells, respectively for 1 hr at 37°C. Next the mixtures were removed from cells, replaced with fresh complete DMEM medium, and incubated at 37°C for 24 hr and 72 hr respectively. At indicated times, cells were harvested for quantification of GFP-positive cells by flow cytometry using BD Accuri C6 flow cytometer (BD Biosciences). “Neutralizing activity” represents the number of GFP^+^ cells detected after incubation of virus in AW from IL-13 or vehicle treated HAE.

### Virus

The 2019n-CoV/USA-WA1/2019 (WT-SARS-CoV-2) was kindly provided by BEI Resources. SARS2-N501Y_MA30_, a mouse adapted SARS-CoV-2 virus was generated by serial passage through mouse lungs 30 times as reported previously (31). Two replication-competent GFP-expressing recombinant viruses were used: Sendai virus-eGFP (SeV-eGFP), a gift from Dr. Ultan Power (24) and VSV-eGFP-SARS-CoV-2, a gift from Dr. Wendy Maury. These two recombinant viruses were prepared as described in previous reports (24, 61–63).

### Virus titration

Viral loads in tissues were measured by plaque assay using VeroE6 cells. In brief, tissues were homogenized in DMEM with one freeze-thaw cycle for releasing intracellular virions. 100 μL homogenized tissues were serially diluted with 900μL DMEM, i.e., 10^-1^, 10^-2^, 10^-3^, 10^-4^, 10^-5^. We added 200μL of the sample dilutions to designated 12-wells and incubated at 37°C for 1 hr. After removing inocula, 1.2% agarose and 2X DMEM media in a 1:1 (V/V) ratio were overlaid on the monolayer cell surface for 3 days of incubation. On third day, the ovverlay was removed and virus-induced plaques were visualized by 0.1% crystal violet staining. Viral titers were expressed as the number of plaque-forming units (PFU) per mL tissue.

### Immunofluorescence

Primary cultured HAE were washed with PBS (Thermo Fisher Scientific), fixed with 4% paraformaldehyde (Electron Microscopy Sciences) in PBS for 15 minutes, permeabilized with 0.3% Triton-X (Thermo Fisher Scientific) in PBS for 20 minutes and then blocked with 2%BSA (Research Products International) in Superblock (Thermo Fisher Scientific) for 1 hour at room temperature. Next, the cultures were incubated with primary antibody apically for 1.5 hours at 37°C, then incubated with secondary antibody apically for 1 hour at 37 °C. Both primary and secondary antibody were diluted in 2% BSA and Superblock. Samples were labeled with following primary antibodies: mouse anti-MUC5AC (1:2000, Thermo Fisher Scientific #MA5-12178), anti-N protein (1:1000, CiteAb #40588-T62) and mouse anti-acetylated α-tubulin (1:200, Sigma-Aldrich #T7451). The secondary antibodies used were goat anti-mouse and goat anti-rabbit conjugated to Alexa Fluor 568 or 488 (1:500, Thermo Fisher Scientific #A-11004, #11008). The cultures were washed with PBS, mounted in Vectashield with DAPI (Vector Laboratories) and cover slipped.

### Histopathology and Immunohistochemistry

Tissues were placed in 10% neutral buffered formalin (3-5 days), dehydrated through a progressive series of alcohol and xylene baths, paraffin-embedded, sectioned (∼4 µm) and stained with hematoxylin and eosin (HE) stains for general examination or diastase-pretreated periodic Schiff (dPAS) stain for mucus detection (64). Tissues were examined by a veterinary pathologist using the post-examination masking technique (65). Healthy mouse lung lacks airway goblet cells in airways. To test the effects of IL-13 we ordinally scored cross-sectioned airways for the severity of Goblet (mucous) cell changes: 0 – none; 1 – scattered; 2 – circumferential around airway; 3 - Circumferential and partial mucus accumulation in lumen; and 4 - Circumferential and overt mucus obstruction of airway. Acute lung injury was defined by the extent of lesions (edema/hyaline membranes) consistent with the effusive phase of diffuse alveolar disease: 0, absent; 1, 0-25%; 2, 26-50%; 3, 51-75% and 4, >75% of tissue fields. Immunohistochemistry for SARS-CoV-2 N protein was conducted as previously done with minor optimizations (31). Briefly, paraffin-embedded sections were treated for epitope retrieval (citrate buffer, pH 6.0, 110°C for 15 minutes) followed by primary antibody (1:4000 x 15 minutes, rabbit polyclonal, Sino Bio 40143-T62), secondary antibody kit (DAKO Rabbit Envision HRP System), and chromogen (DAKO DAB Plus). Ordinal scoring for lung N protein was performed as previously: 0 – absent, 1 – 0-25%; 2 – 26-50%; 3 – 51-75% and 4 - >75% of tissue fields.

### Single-cell RNA-seq Library preparation

Cells were fixed, permeabilized and 8 cDNA sub-libraries were generated using the SPLiT-seq (66) protocol V3.0 from Parse Biosciences following the manufacturer’s instructions. Concentration and fragment size of the pooled cDNA library was measured using Qubit HS and High Sensitivity DNA Assay. Fragment size for the library was ∼430bp which falls within the recommended 150-850bp.

### Single-cell RNA Sequencing

Pair-end reads sequencing was performed on the pooled library using two lanes of Illumina NovaSeq 6000 S1 Flowcell instrument (approximately 800 million PE reads per lane) for 300 cycles to generate 150bp paired-end reads. The sequencing was configured for 66nt (transcript sequence) read1, 94nt (cell-specific barcodes and UMI) read2, and an additional 6nt read1 index for sub-library indices as described in the manufacturer’s protocol. Sequencing was performed by the University of Iowa Genomics Facility.

### Single-cell RNA-seq Data processing

Human- SARS-CoV-2 combined reference genome was made using human reference genome-build GRCh38.p13 (accession GCF_000001405.39) and SARS-CoV-2/human/USA/WA-CDC-WA1/2020 isolate (GenBank assembly accession: GCA_009937905.1). Pair-ends reads obtained from Illumina NovaSeq were processed and mapped to the human-SARS-CoV-2 combined reference genome using STAR and Samtools package implemented via Split Bio data analysis pipeline; split-pipe v0.7.5p in Amazon Web Services platform according to the suggested data processing protocol by the manufacturer. Output files (a matrix with cell-gene counts, cell metadata file, genes info file) from all 8 sub-libraries were imported and merged into a single Seurat object using Seurat v4.0.1 in R v4.0.5 for further analyses. Low-quality and duplicated cells were filtered and only cells with 1) % of mitochondrial genes <0.5, 2) 2500<UMI counts per cells >40,000, 3) 1,000 < number of genes per cells >8,000 was selected for downstream analysis (**Supplementary FigureS1**).

### Identifying Cell types in Single-cell RNA-seq data

Cell types were identified using the combination of 1) cells clusters identified by Seurat’s FindClusters function (uses shared nearest neighbor modularity optimization based clustering algorithm; identified 19 clusters using resolution 0.6), 2) expression of airway epithelial cell-type specific markers: Basal (KRT5, TP63, KRT6A), Ciliated (FOXJ1, TPPP3), Secretory (SCGB3A1, BPIFA1), Goblet (MUC5AC, LYZ, ADRA2A, PRB2, FCGBP), Ionocytes(FOXI1, ASCL3), Mitotic (MKI67, TOP2A), Pulmonary Neuro Endocrine cells (CALCA, ASCL1), Intermediate (clusters expression basal, ciliated and secretory cell markers), 3) Clusters expressing greater than 2 fold of specific cell type markers (adjusted p-value<0.05) in multiple pairwise differential expression analyses using Seurat’s FindMarkers function(uses Wilcoxon rank sum test). Since, not all goblet cells clustered together in a specific cluster, to further expand the number of goblet cells, the sum of the log normalized counts for goblet cells specific markers (MUC5AC, LYZ, ADRA2A, PRB2, FCGBP) was calculated. Cells were labelled as goblet if 1) Cells were present in clusters 1,13 or 16 (containing majority of secretory cells), 2) sum of normalized goblet cell marker genes were greater than 2 (**Supplementary FigureS2**).

### Identifying SARS-CoV-2 infected cells and analysis of background contamination in combinatorial single cell RNA-seq data

Whereas single-cell RNA-seq can accurately quantify transcripts, it may also be affected by background levels of transcripts present outside of cells during the RNA-seq library preparation; it is particularly important to control for background contamination when quantifying transcripts that may be highly abundant in some cells but not others.

To estimate background contamination using empirical data, we took advantage of our dataset containing sequencing data from primary airway epithelial cells from both male and female humans, and from samples exposed vs. unexposed to SARS-CoV-2. First, we calculated % of misassigned UMIs per cell using Y chromosome genes (DDX3Y, EIF1AY, KDM5D, RPS4Y1, TTTY15, UTY, ZFY, NLGN4Y, LINC00278, LOC107987344) and donor information (Donor C: male, Donor A/B: female); if a “cell” from female sample showed high expression of Y chromosome genes, it was labelled as misassigned. Then using SARS-CoV-2 viral transcripts genes (orf1ab, orf7a, orf8, orf3a, orf10, orf6, E, M, N, S) and viral exposure information, we calculated % of misassigned UMIs per cell; if a cell from group unexposed to SARS-CoV-2 showed expression of SARS-CoV-2 viral transcripts, the UMI was labelled as misassigned. We then plotted % of misassigned UMI per cell for each Y chromosome and viral transcripts genes against total gene counts in our data. Interestingly, spurious detection of viral transcripts may occur at higher rates than for host genes, however, comparison of (total UMI count vs. genes) shows the majority is occurring in low depth cells, etc., which allowed us to establish a threshold for filtering cells (**Supplementary FigureS3, a and b**). We then used stringent criteria based on clustering and on our background contamination analysis to define a cell as “infected”. We first calculated the sum of SARS-CoV-2 transcripts (orf1ab, orf7a, orf8, orf3a, orf10, orf6, E, M, N, S) using normalized counts for each cell and applied the following criteria: 1) belongs to the cluster with high counts of SARS-CoV-2 transcripts as identified by UMAP, 2) belongs to a sample infected with SARS-CoV-2 virus, and 3) sum of SARS-CoV-2 transcript normalized count is greater than 3 (**Supplementary FigureS3, c-f)**.

### Mixed model DEG analysis

Differential expression analysis was performed using a generalized linear mixed model. The gene count for each cell was assumed to follow a negative binomial distribution. The log of the expected count was a linear function with fixed effects for treatment and donor, where donor was included to account for pairing of treatments from the same donor, and a random effect for sample, to account for batch effects. The log of the total number of unique molecular identifiers (UMIs) for each cell was used as an offset to account for different depth of information between cells. The R package lme4 (67) was used to fit the model, and log2 fold change and p-value for each gene were the estimated coefficient and Wald test p-value for treatment, respectively. After testing all genes, adjusted p-values were obtained using the Benjamini-Hochberg criterion to control false discovery rate (FDR) (68).

### Bulk RNA-Sequencing

Total RNA was extracted and purified from harvested mice lungs using RNeasy plus mini kit from QIAGEN. cDNA library was prepared using standard Illumina TruSeq stranded mRNA protocol. Concentration and fragment size of the library was measured using Qubit HS and High Sensitivity DNA Assay. The pooled library was sequenced on one lane of Illumina NovaSeq 6000 SP Flowcell instrument for paired end reads. Library preparation, quality control and sequencing were the done by the University of Iowa Genomics Facility. Base cells were demultiplexed and converted to FASTQ format by the sequencing core. The processed fastq files were pseudo aligned to mouse reference (69). Mapped raw reads were reported in transcripts per million (TPM) by Kallisto. For downstream analysis, output from Kallisto was summarized into gene-level estimates using tximport v1.10.1 in R v3.5.1 (70). Differential gene expression was performed using iDEP v0.92 (71) and DESeq2 v1.30 (72). Pathway analysis was done using fast gene set enrichment analysis (fgsea) and msigdbr v7.5.1 (73) packages in R v3.5.1.

### Ingenuity Pathway Analysis – Upstream regulator analysis

Upstream regulator analysis was done using QIAGEN IPA (74) with the confidence level set to experimentally observed only. Genes included passed a threshold of an adjusted P value of less than 0.1 and a log2 absolute fold change of greater than 1. Upstream regulators were sorted according to the predicted activation Z score.

### Mouse model

8-9-month C57BL/6J mice and 6-10-week BALB/c mice (n=4) were intranasally infected by 1,000PFU of SARS2-N501Y_MA30_ after a 4-day pretreatment of 2.5 μg/day intranasal IL-13 or vehicle (21). 6-10-week (young) BALB/c mice and 8-9-month (middle-aged) C57BL/6J mice were acquired from the Jackson Laboratory. The *Pla2g2d*^-/-^ mouse strain was previously reported (40). Mice were maintained in the Animal Care Unit at the University of Iowa under standard dark/light cycle conditions, ambient temperature, and humidity.

### LC-MS/MS

Mouse lungs were harvested and homogenized into 1mL methanol/lung tissue using a tissue homogenizer with beads. After centrifugation at 16,000 rpm for 10 min at 4°C, 200 μL supernatant was transferred to a clean tube and diluted with HPLC grade water to a final volume of 2 mL of 10% methanol (V/V). 0.1 ng/μL Internal standard cocktail was added in diluted samples and serially diluted primary standards (4.76, 0.476, 0.0476, 0.00476, 0.000476 ng/μL). Then, samples and primary standards were subjected to solid phase extraction using Strata-X 33 μm Polymeric Reversed Phase columns (60 mg/3 mL). After pre-washing columns with 3 mL methanol and water, each sample was loaded on the column, and lipid extracts were eluted with 1 mL methanol, vacuum dried, and suspended in 100 μL of mobile phase A. Ten eicosanoids were separated on an Acquity UPLC BEH C18 column using series dilution with mobile phase A and phase B at a 300 μL/min flow rate. Each eicosanoid was identified by the retention time of its corresponding primary standard.

### Statistics

Statistical analysis was performed using Student’s unpaired or paired 2-tailed t test or log-rank (Mantel-Cox) test or one-way ANOVA test. For RNA-seq data, p-values were corrected using Benjamini-Hochberg FDR. For scRNA-seq data, differential gene expression was performed using Wilcoxon rank sum test with Bonferroni correction in Seurat. For bulk RNA-seq, differential gene expression was performed using DESeq2 in iDEP v0.92. Data are represented as mean or mean ± SEM or mean ± SD. P-value < 0.05 was considered statistically significant.

### Ethics statement

Primary airway epithelia cells were obtained from the University of Iowa Cells and Tissue Core. Mouse studies were reviewed and approved by the Institutional Animal Care and use Committee of the University of Iowa following the protocol #2071795. Mice were maintained in the Animal Care Unit at the University of Iowa under standard dark/light cycle conditions, ambient temperature, and humidity.

### Data availability

Human scRNA-seq and mouse bulk RNA-seq data has been deposited in the NCBI’s Gene Expression Omnibus (GEO) database (GSE249308). Supporting data values are provided as a supplementary table with the paper. Additional data are available from the corresponding author, following publication upon reasonable request.

### Author contributions

Research study design (S.G., B.X., K.L., C.L.W, A.L.T, D.K.M, S.P., P.B.M., A.A.P.). Conducting experiments and acquiring data (S.G., B.X., K.L., R.M.G., C.L.W., A.L.T., H.G., G.G., J.Z.). Analyzing data (S.G., B.X., K.L., R.G., A.L.T., G.N., D.K.M., S.P., P.B.M., A.A.P.). Manuscript draft review (all authors), writing the manuscript (S.G., B.X., P.B.M., A.A.P.). Co-first authorship order was assigned based on temporal involvement in the project.

## Acknowledgments

This study was supported by Carver Trust COVID-19 Grant and CF Foundation Iowa RDP (A.A.P. and P.B.M). A.A.P was supported by NIH 1R01HL163024 and K01HL140261, Cystic Fibrosis Foundation PEZZUL20A1-KB and the Stead Family Foundation. P.B.M. was supported by NIH R01AI129269, the Cystic Fibrosis Foundation and the Roy J. Carver Charitable Trust. P.B.M. and S.P. were supported by NIH P01AI060699. The Carver College of Medicine Cell and Tissue and Comparative Pathology cores are in part supported by the Center for Gene Therapy for Cystic Fibrosis (NIH Grant P30 DK-54759). We acknowledge the support from Cells and Tissue Core, Comparative Pathology Core, Genomics facility and Animal Care unit of University of Iowa as well as Charlie Roco from Parse Biosciences. We also thank BEI Resources, Ultan Power and Wendy Maury for providing SARS-CoV-2 and recombinant viruses.

