## Supplementary Figures for "IL-13 decreases susceptibility to airway epithelial SARS-CoV-2 infection but increases disease severity in vivo"

**SUPPLEMENTARY FIGURE AND LEGENDS**

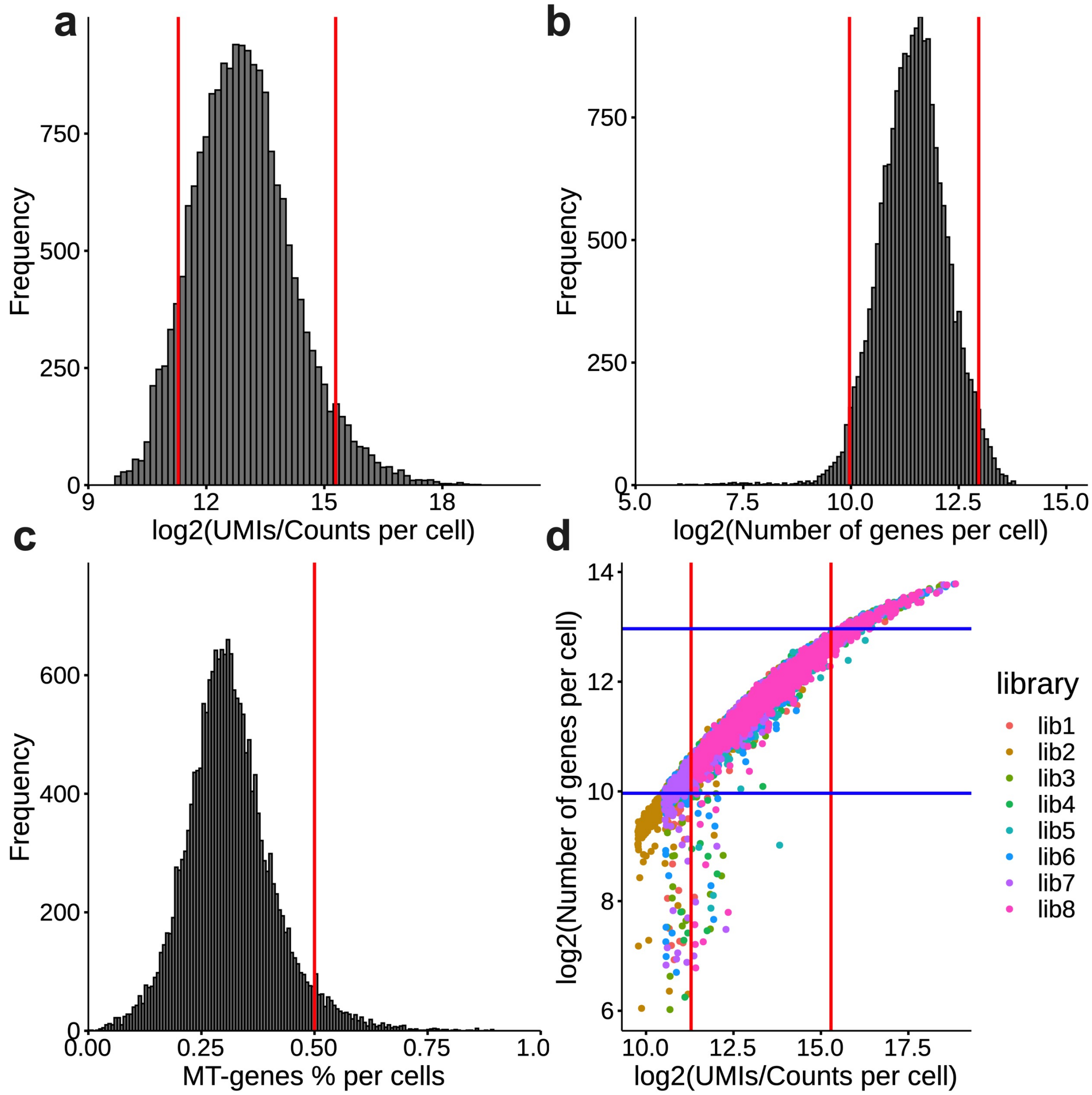


**Supplementary FigureS1. scRNA seq data QC.** Figure shows (a) the histogram of log2 (UMIs/Counts per cell), (b) log2 (number of genes per cell), (c) % of Mito genes per cell and (d) scatter plot of UMIs vs Number of genes detected per cell in log2 scale, colored by each sub-library. Red lines in each graph represents the cutoff (% of mitochondrial genes <0.5, 2,500 < UMI counts/cells > 40,000, 1,000 < Number of genes/cells > 8,000). Figure D shows cutoff for both A and B.

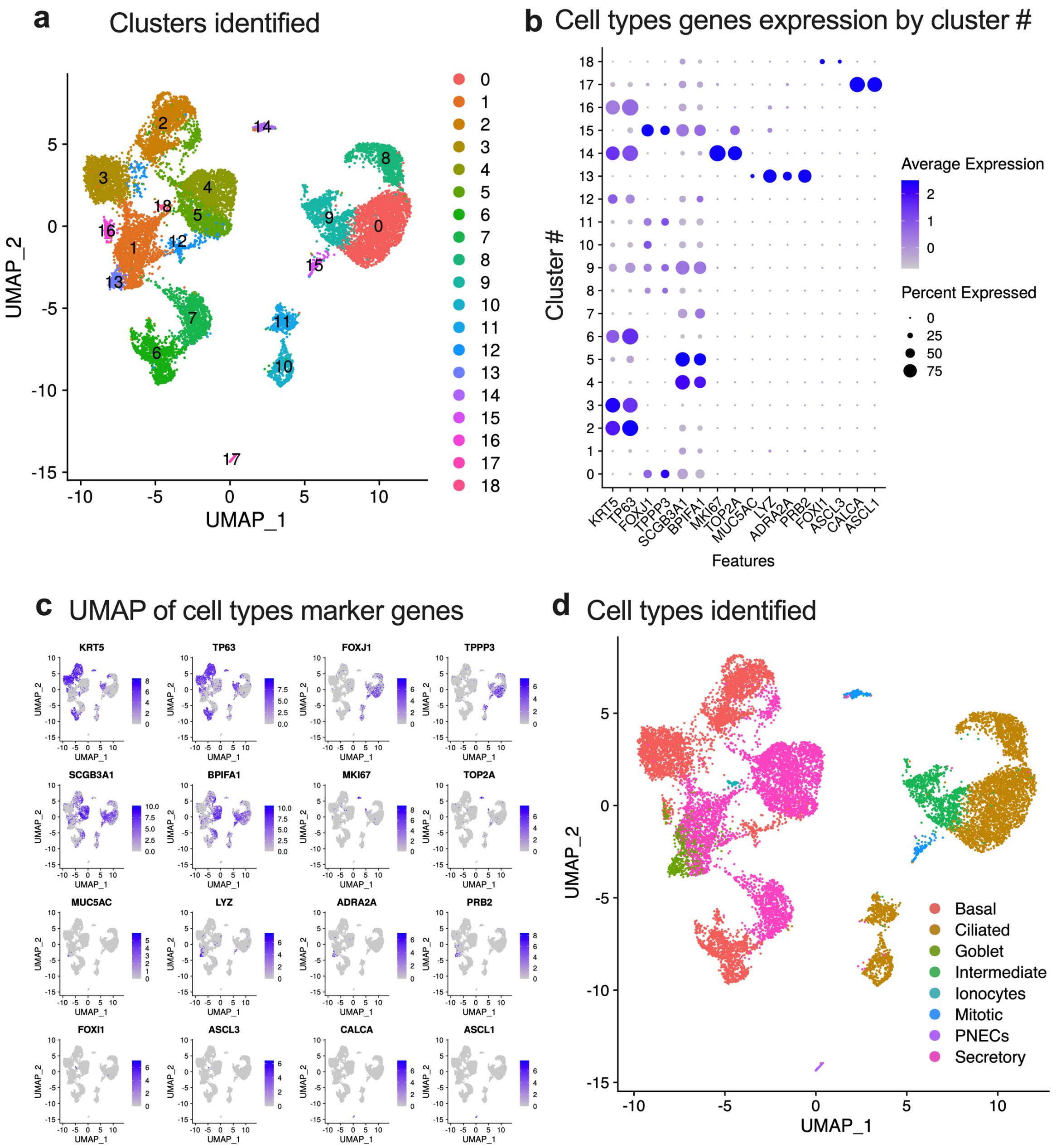


**Supplementary FigureS2. Identifying cell types**. (a) Cell clusters identified by Seurat's FindCluster function using resolution 0.6. (b) Dotplot showing the expression of cell type specific gene in identified clusters. (c) UMAP of cell type marker genes (d) UMAP of identified cell types in the scRNA seq data.

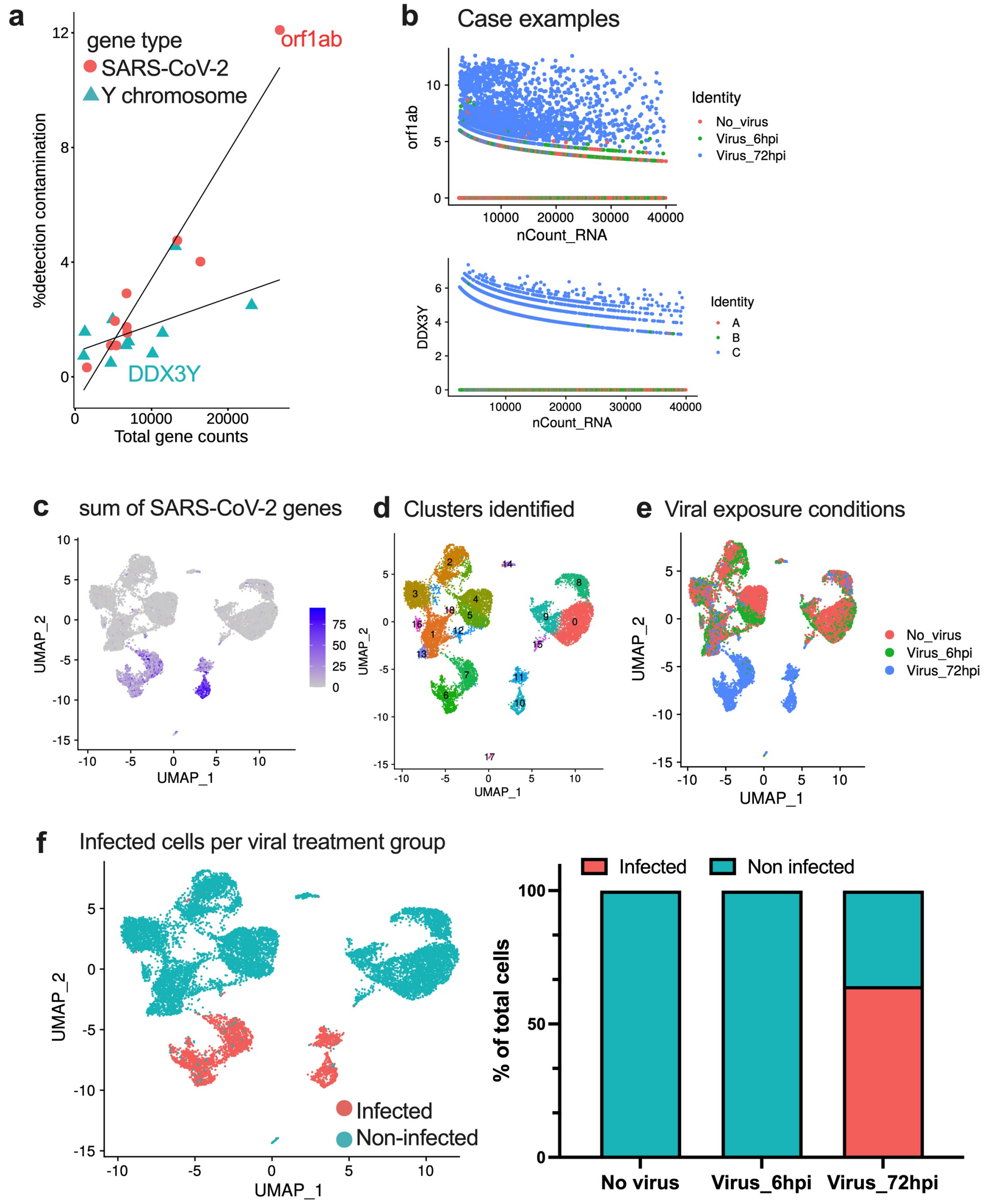


**Supplementary FigureS3. Detection of suspected background contamination and Identifying SARS-CoV-2 infected cells.** (a) Correlation plot showing % of misassigned UMIs VS total gene counts for each gene in the dataset, (b) Case examples showing the expression level of misassigned genes. (c-e) UMAPs showing the sum of SARS-CoV-2 transcripts, clusters identified and viral treatment groups respectively. Combination of (c-e) was used to identify infected cells shown in (f). (f)-right shows the proportion of infected cells identified in each viral treatment group.

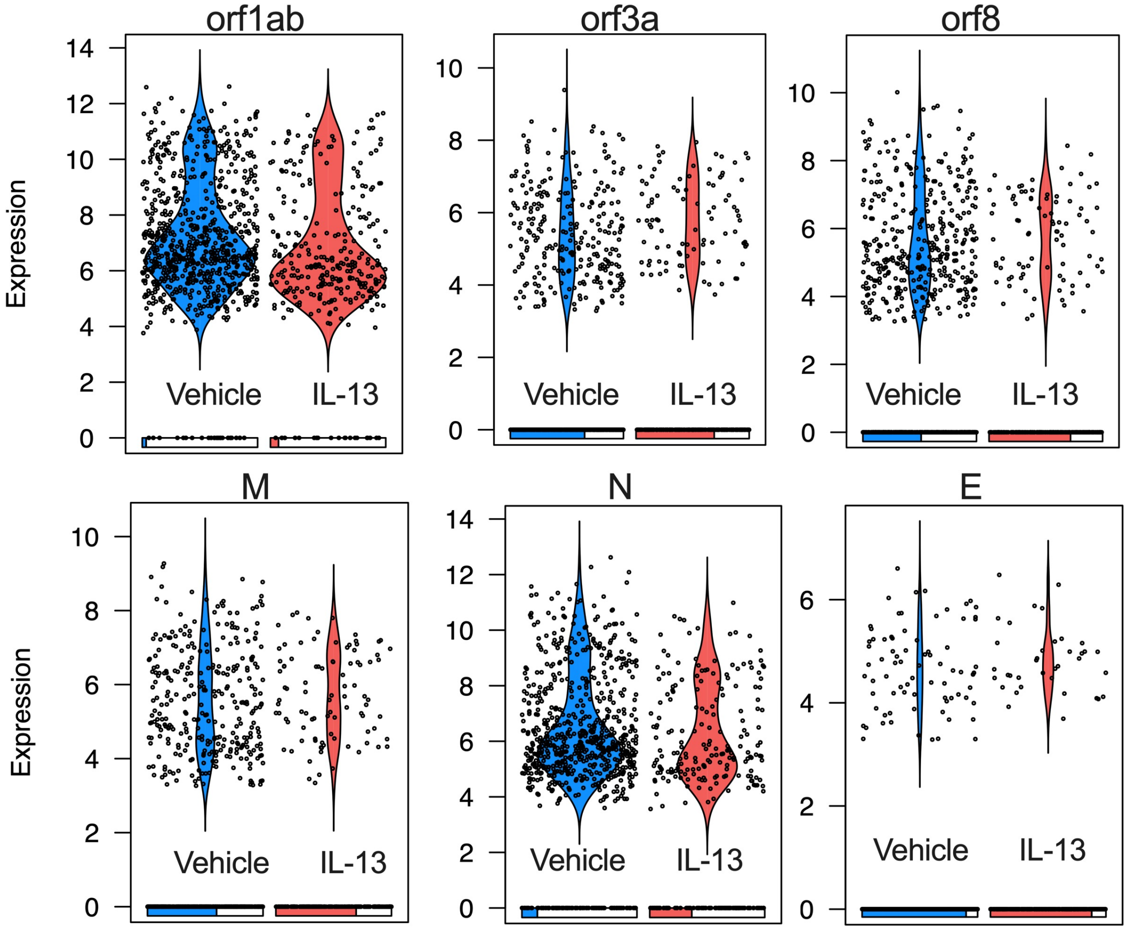


**Supplementary FigureS4**. **Expression of viral genes in infected cells.** Violin plot of SARS-CoV-2 transcripts in Vehicle vs IL-13 (55days) treated HAE at 72hpi with SARS-CoV-2. Each dot represents a SARS-CoV-2 transcript-positive cell. The bar at the bottom shows the proportion of cells with zero expression in each group.

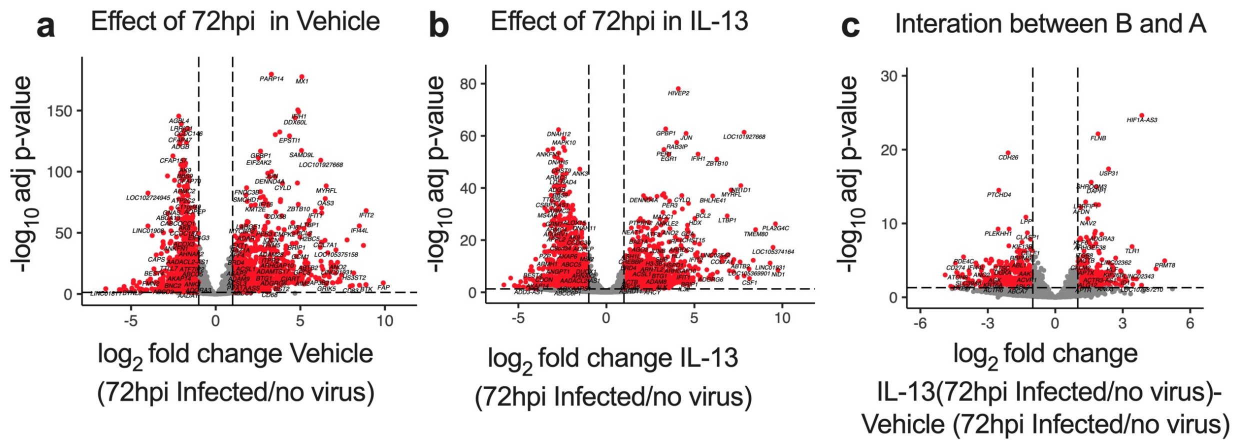


**Supplementary FigureS5. Effect of IL-13 treatment on the response of infected ciliated cells.** Volcano plots showing DEGs induced by SARS-CoV-2 72hpi in infected ciliated cells from (a) Vehicle and (b) IL-13 (20ng/mL, 4days) treated HAE. (c) shows the interaction between (a) and (b). Threshold used for calling significant genes (labelled by red dots): abs ($log_2$fold change)>1, adj p-value<=0.05.

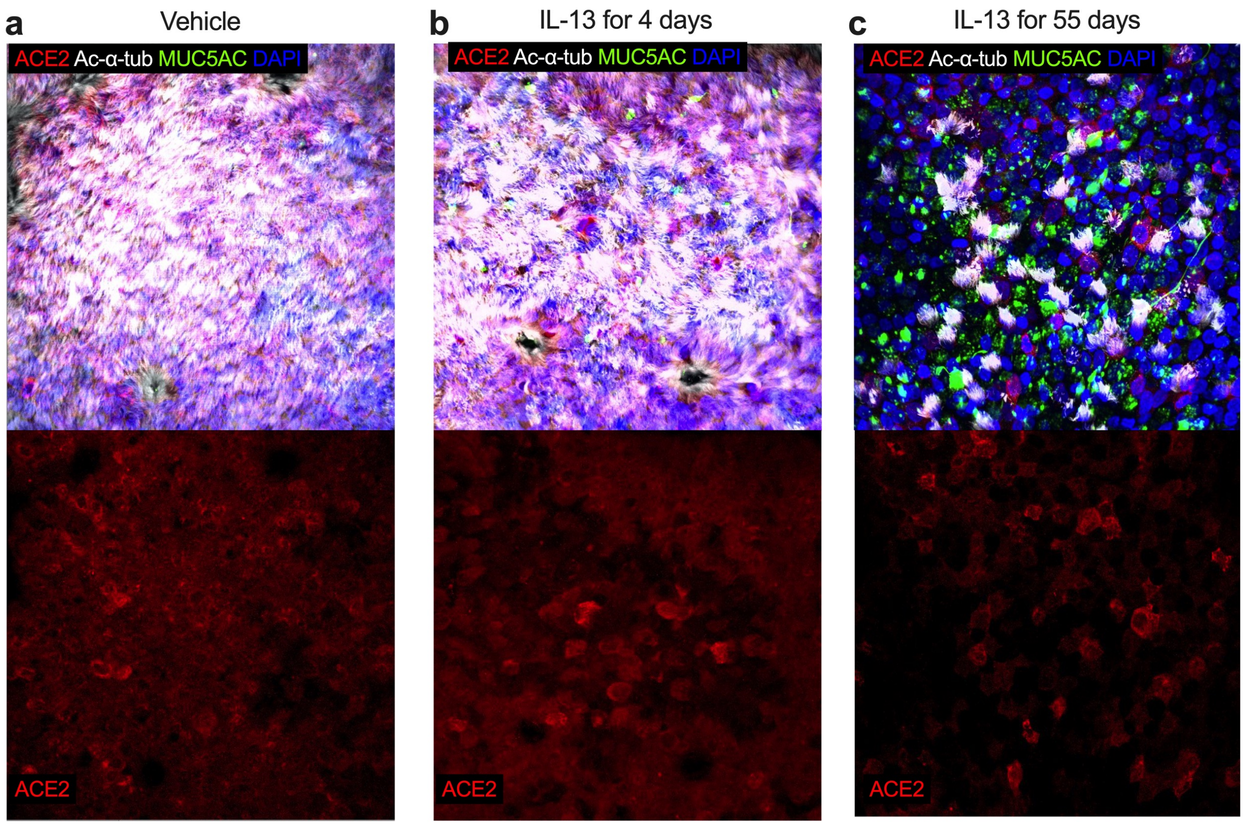


**Supplementary FigureS6. IL-13 decreases abundance of ACE2-expressing ciliated cells.** Immunofluorescence confocal microscopy for ACE2 (red, bottom panel), acetylated alpha tubulin (ciliated cells, white), MUC5AC (goblet cells, green) and DAPI (nuclei, blue).

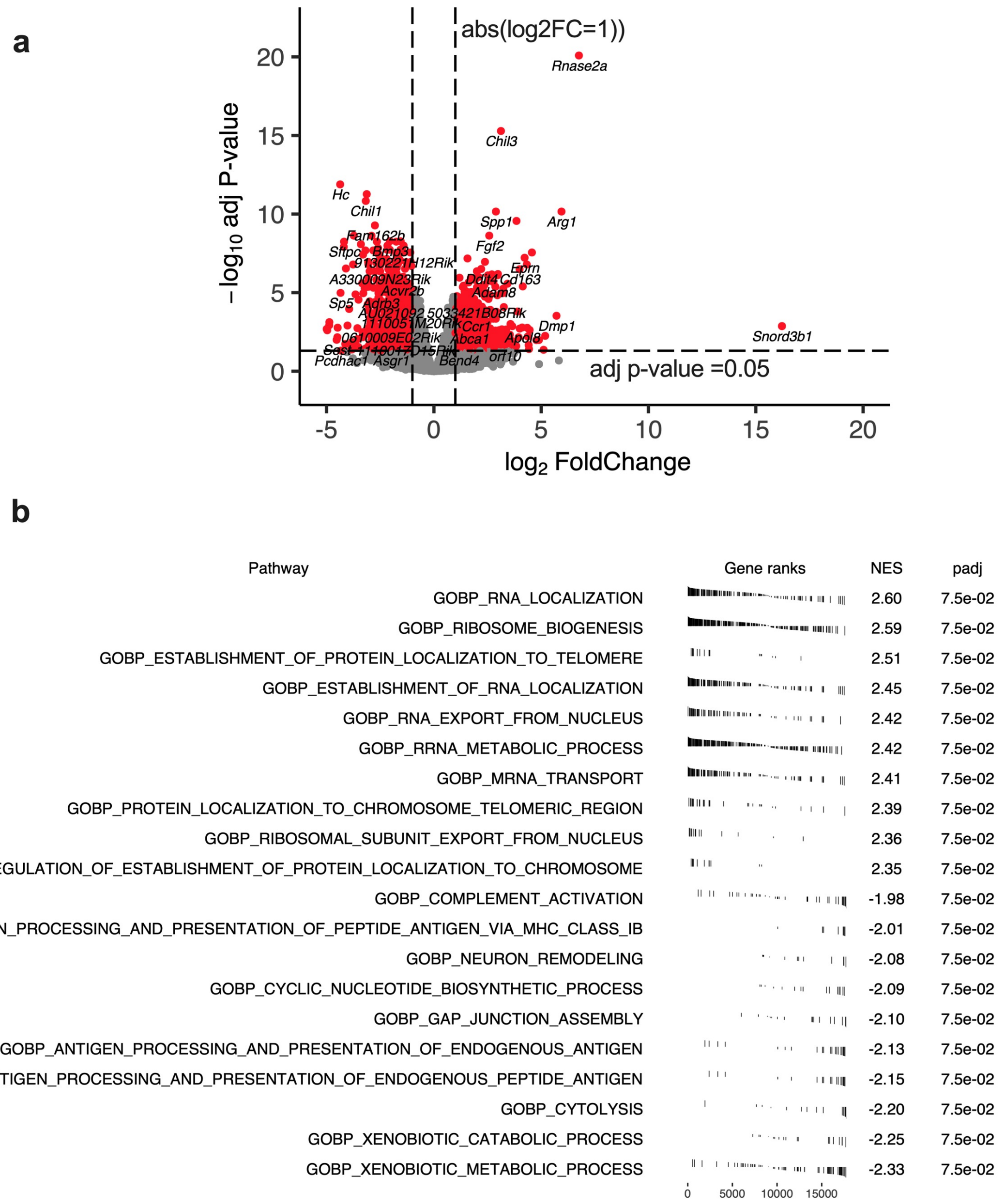


**Supplementary FigureS7**. **Pathway analysis of infected mice pre-treated with Vehicle or IL-13 (2.5 µg/day intranasal IL-13) for 4 days.** (a) Volcano plot showing DEGs between Vehicle and IL-13 groups. Threshold for significant genes (red): abs (log2foldchange) > 1, adj p-value < 0.05 is shown by dotted line. (b) Gene set enrichment analysis (GSEA) table from pathway analysis showing normalized enrichment score and adjusted p-value. Pathway analysis was done using GO: Biological process database.

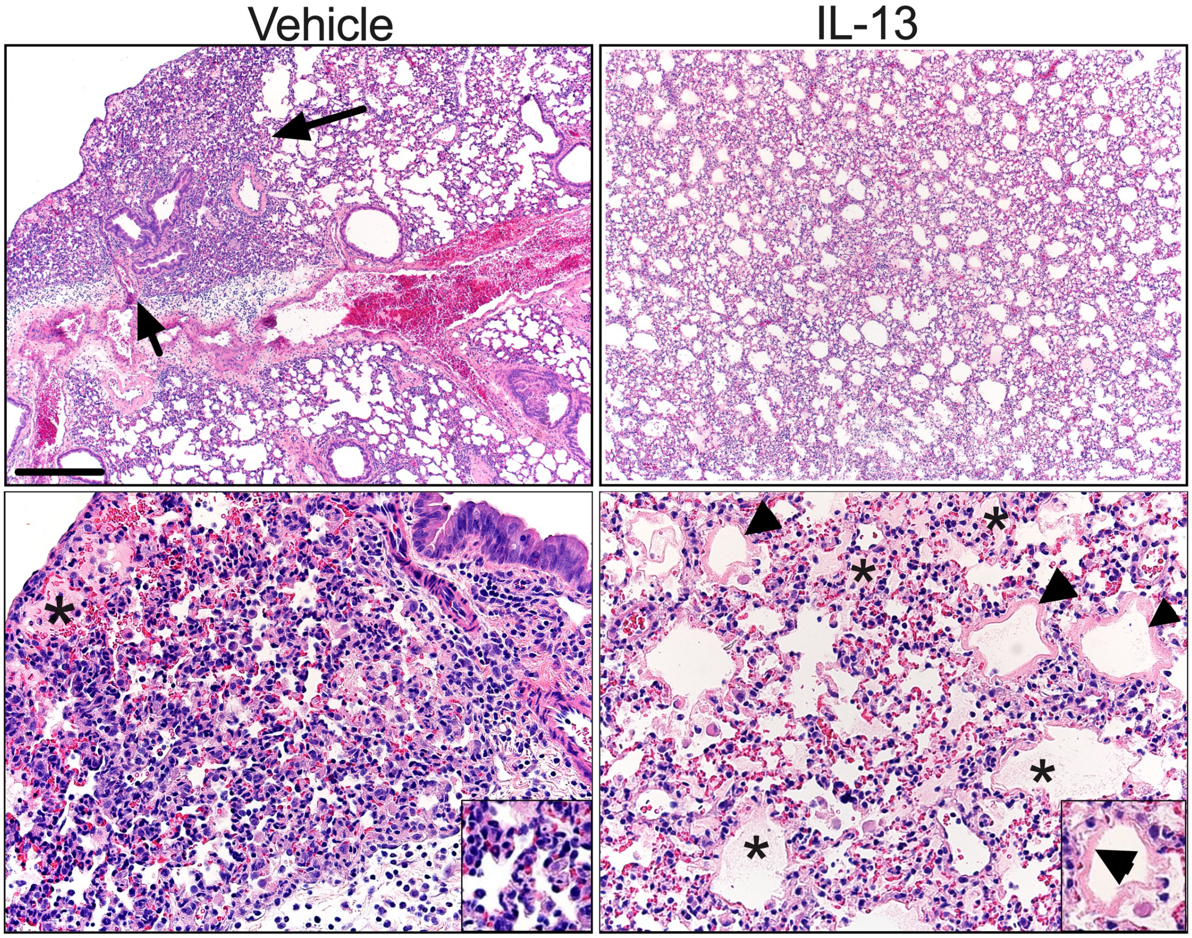


**Supplementary FigureS8. IL-13 increases SARS-CoV-2 -induced edema and hyaline membranes.** Veh or IL-13 treated samples (left and right columns respectively) at 5 d.p.i. of SARS-CoV-2 infection. Veh treated group samples were distinguishable by regions of consolidation (arrows, and inset, left column) composed of atelectasis, increased cellularity by leukocytes, cellular, debris and localized edema (*). In contrast, IL-13 treated samples lacked regions of consolidation and had more evidence of edema (*) and hyaline membranes (arrowheads and inset, right column). HE stains, bar = 392 and 78 µm, top and bottom rows respectively).

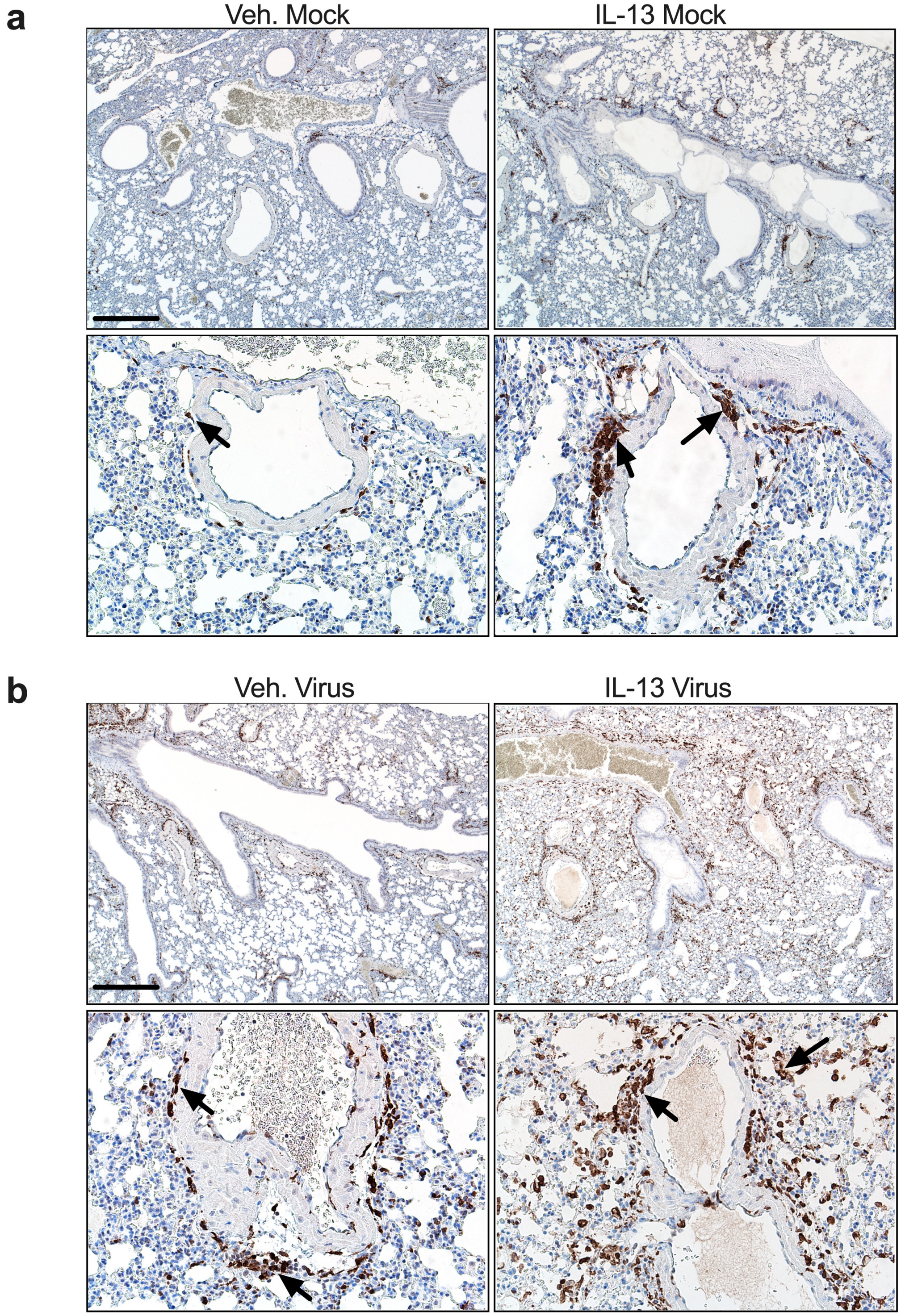


**Supplementary FigureS9. IL-13 increases SARS-CoV-2- induced peri-vascular M2 macrophages.** Vehicle or IL-13 treated sample with (a) mock or (bb) 5dpi SARS-CoV-2 infection. IL-13 treated samples had more AIF1/IBA1 brown staining (arrows) compared to vehicle on low mag of lung and surrounding pulmonary veins.
Top bar = 425 μm and bottom bar =85 μm.
